# Multiple-serotype models of dengue virus transmission: simulation study and perspectives for the application of inference in epidemiological surveillance

**DOI:** 10.1101/583351

**Authors:** Caetano Souto-Maior

**Author notes:** current address: National Heart Lung and Blood Institute, NIH, Bethesda, MD, United States of America.

## Abstract

With around 3 billion people at risk, dengue virus is endemic to many parts of the world. In the Brazilian city of Rio de Janeiro, surveillance measures require notification of new dengue virus cases, and are supplemented by serum collection from patients and sequencing of viral RNA. Phylogenetic analyses have been performed for all serotypes circulating in the country to identify viral genotypes, potentially identify new introductions, and compare viruses presently circulating in the country with those in the past, and of other countries. As a separate type of analysis, a number of mathematical models have been developed to describe dengue virus transmission – particularly qualitative incidence or prevalence patterns – although few have been tested. In this chapter, I show how different mathematical formulations could represent transmission of dengue virus by mosquitoes to humans, how the different model structures entail assumptions about the process, and how these affect outputs qualitatively. Inference from simulated data is used as proof of principle that the kind of data available could be used to accurately estimate all model parameters; however, it is shown that stochasticity may severely hamper efforts to test the models quantitatively. I further implement inference from sequence data for the different models, and compare the performance to that of time series. The methods are applied to the data available for the city of Rio de Janeiro.

## 1 Background

The persistence of dengue fever (as well as more severe syndromes caused by dengue virus) constitutes the most extensive viral epidemic transmitted by arthropods, with around 3 billion people at risk worldwide, and 300 million annual cases estimated (Bhatt et al. 2013). The recently recorded expansion in the range of the main transmission vectors, *Aedes aegypti* and *Aedes albopictus* (Kraemer et al. 2015) *–* presumably due to higher temperatures at temperate regions resulting from climate change – in combination with the emergence of other *Aedes*-transmitted diseases further increased attention to vector control.

Although among the vector transmitted viruses dengue itself has arguably lost some of the attention to Chikungunya and especially to Zika virus (due especially to the previously unknown relationship between the latter and birth defects) at the population level the study of either of these diseases is to a very large extent the study its human and mosquito hosts. The fact that licensed vaccines for these disease was essentially absent – dengue virus had a vaccine in phase 3 clinical trials (Eisen & Moore 2013; Hadinegoro et al. 2015; Villar et al. 2014) that was just recently licensed (WHO 2016) – further highlights the importance of vector control, and of the knowledge about dengue virus transmission in the control of all of the diseases transmitted by the *Aedes* mosquitoes.

One half of the of cycle of dengue virus (*DENV*) – that is, the mosquito-to-human transmission – happens through the bite of an infected *Aedes aegypti* or *Aedes albopictus* mosquito (i.e. the vector in *vector-transmitted disease*); the transmission cycle is completed when an infected human is bitten by a mosquito that in turn becomes infected (Halstead 2007). Although importation from other geographical areas (Salje et al. 2012; Stoddard et al. 2013; Vazquez-Prokopec et al. 2010) as well as sylvatic cycles between non-human primates and other *Aedes* species may play a role in sustaining transmission (Vasilakis et al. 2011), these basic steps of the human-*Aedes* cycle should be enough to create chains of transmission that allow endemicity, and this is considered the primary cycle in explaining dengue virus persistence.

A few details are noteworthy in a general model of dengue virus transmission, which would otherwise conform nicely to that of a generic vector-transmitted mode of propagation. Dengue virus has four antigenically distinct variants – types, or strains – commonly referred to as serotypes (DENV-1 through 4). Anything from a single one to all four of them can be circulating in any one place. If only one serotype is present, a simple description of transmission where a susceptible host gets infected, recovers, and becomes immune to further infection is generally adequate, since infection with a serotype is accepted to confer human hosts lifelong immunity against that same type. If more than one serotype is circulating, infection can happen at least twice (but not more than four times, because unlike influenza, for instance, evolution of the virus does not allow it to escape immunity built against it), one for each previously unseen serotype. In this case multiple infections may need to be accounted for. Also, it could be important to differentiate between strains, as a secondary infection can only be caused by a serotype different from the previous.

Dengue virus transmission has been extensively explored through mathematical models (Johansson et al. 2011). As usual, the disease states of the human hosts have been described by simple extensions of the susceptible-infected-recovered framework, often (but not always) coupled with susceptible-infected description of the mosquito hosts. A mix-and-match of other known or suspected features specific (although possibly not exclusive) to dengue virus have been further added: secondary infections, temporary strain-transcending immunity (cross-protection), enhanced (or reduced) susceptibility to secondary infections, increased lethality in case of severe presentations (often associated to secondary infections, as well as other risk factors such as age or blood-related dysfunctions) (Johansson et al. 2011).

Therefore, dengue virus transmission can be described mathematically by multiple explicit serotypes, which we denote by multiple-serotype models (e.g. a two-serotype model means two different strains of dengue are explicitly described), or by a single explicit virus type, hereafter denoted by SIRX models (which include the classic SIR and SIRS models, as well as intermediate formulations, as described in the methods section). Both formulations purport to describe settings where one or more serotypes may be circulating; in the former description each explicit serotype causes infection once, while in the latter the ensemble of unspecified serotypes causes infection twice or more.

Although seemingly subtle, the conceptual difference between the two modeling approaches is profound: while in the SIRX vector models secondary infections depended exclusively on waiting for recovered individuals to become susceptible again, in the multi-type models strains compete for multiply susceptible individuals. This feature can causes serotype alternation and induce oscillation even in the absence of seasonal forcing. More importantly, this model is less of a caricature of the process of disease transmission, since it is widely accepted that an individual infected with a serotype cannot be infected again by the same strain, rendering the explicit description of the SIRX models technically impossible. Whether one approach or the other is more suitable to describe real dengue epidemics, however, cannot in principle be decided without confronting both models to epidemiological data.

On the epidemiological records side of dengue, some unique patterns are often highlighted in dengue virus epidemics, particularly the oscillations with multianual periods and serotype replacement in successive epidemics (Adams et al. 2006). These can be verified, respectively, from incidence records that show greater number of cases usually around the rainy seasons, and through serological surveys or, more recently, sequencing of circulating viruses. Mathematical models of dengue transmission therefore are built to reproduce these broad patterns; nevertheless, different combinations of anyone’s favorite model components may reproduce them, in a way that is indistinguishable from someone else’s choice of building blocks. A non-exhaustive list of processes that could produce realistic outputs in a computer simulation include: stochasticity(Otero & Solari 2010), spatial structure (Favier et al. 2005), enhanced secondary infectivity (Nagao & Koelle 2008), “unnatural” transmission routes (Chikaki & Ishikawa 2009).

One of the most hyped effects among the many incorporated one way or another into the mathematical models is that of antibody dependent enhancement, by which a secondary infection would be more severe than the first in the presence of titers of heterologous antibodies against *DENV* (Kliks et al. 1989). The inclusion of the effect has been shown to drive chaotic dynamics even in deterministic mathematical models (Bianco et al. 2009), so as a result it has been suggested that it could be the most important effect modulating the observed somewhat erratic epidemic patterns. The plausibility of the effect is asserted through the observation that in the presence of subneutralizing antibodies invasion of the cell by viruses is facilitated (Guzman & Vazquez 2010); however, in terms of a mathematical model it is not clear if that would translate into increased susceptibility, increased infectivity, or simply a transmission-unrelated increase in virus lethality. If the magnitude of the enhancement could ever be as large as claimed in modeling studies is not established either. Furthermore, it is not clear whether, if present, the effect would be the dominant factor, or if it would be important to the transmission dynamics at all.

Many other effects and combinations would still not exhaust the list of tens of models that purport to explain dengue transmission (Johansson et al. 2011); nevertheless, a quantitative evaluation of the conformity of these models to real data was not done systematically [but see Rasmussen et al. (2014b) for a rare exception]. Here, I put together a model of the human and vector population with secondary infections, either one or two serotypes, and temporary immunity after any infection, but otherwise minimal in what regards any other asymmetries, enhancements, or alternative routes of transmission. I show that minimalistic multi-serotype models can sustain oscillations in the absence of any of the latter effects or stochasticity; I also implement an individual-based model that can simulate both epidemiological and viral evolution. I further develop inference methods to fit these models to time series and multi-serotype sequence data, compare the inference results for the different kinds of simulated data, and apply the estimation method to real data.

## 2 Methods

### 2.1 SIR model extensions for dengue virus transmission

#### 2.1.1 SIRX plus vector models

The simplest model to describe dengue transmission is arguably the vector SIR model, not unlike the first basic models of malaria transmission with a human and mosquito population, although malaria may have an indefinite number of reinfections, making it a SIS model (Ross 1916; Smith et al. 2012). The SIR model assumes human hosts are only susceptible once (*S*), and after infection (*I*) enter the recovered compartment (*R*) permanently, being immune to any further infection by dengue afterwards.

The vector compartment is modeled as a susceptible mosquito compartment (*U*), and an infected one (*V*), from which a mosquito host never exits once it enters – noting that the female mosquitoes are the only ones transmitting disease, so the male population is absent, or implicit.

Alternatively, human hosts may be allowed to lose immunity acquired from past infections and become susceptible again; under that assumption a single host can potentially be reinfected an unlimited number of times before it dies.

The schematic drawing of both model formulations is shown in figure 1, the only difference between the two being the arrow representing hosts that exit the recovered compartment and reenter the initial susceptible compartment. In this latter case, the structure of the human compartments is that of what is dubbed the SIRS model – the first and last susceptible compartments being the same.

**Figure 1.**
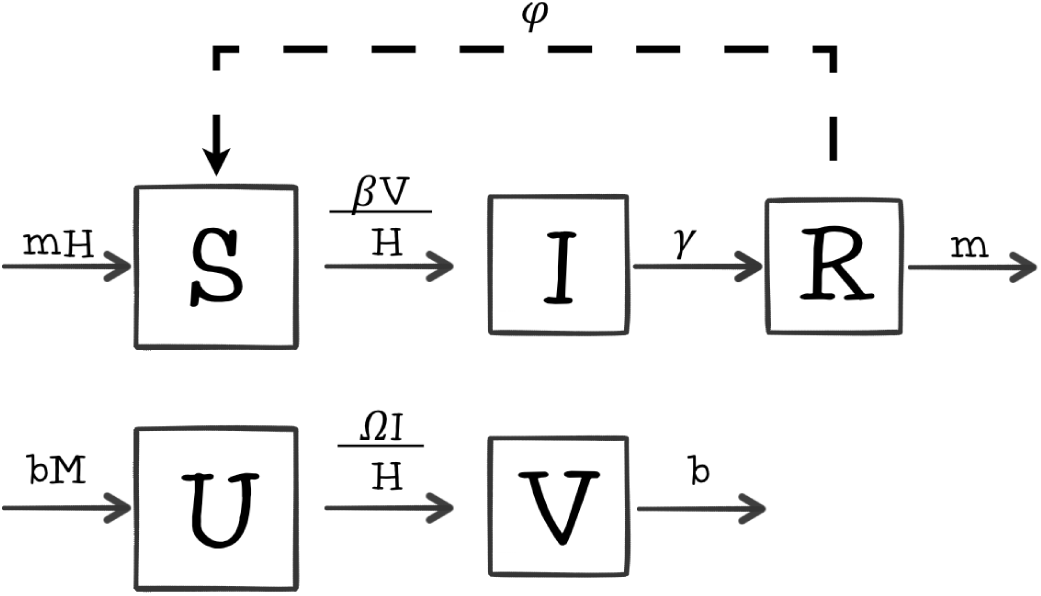
Structure of an SIRS plus vector model, with possible loss of immunity indicated by the dashed arrow. All compartments are subject to natural mortality *m*, but the arrows corresponding to those processes are ommitted in all but the last compartments to avoid repetition and clutter.

The compartments and parameters are very much standard: *m* being the human host mortality rate; *β* the mosquito to human transmission coefficient, or rate; *γ* the human recovery rate; *ϕ* is the immunity loss rate (which is equal to zero in the SIR version of the model); *b* is the mosquito host mortality rate; Ω is the human to mosquito transmission rate. Additionally, it is assumed that the birth rates are the same as the mortality rates for each host species; therefore the population sizes stay constant: *H* is the value set for the human population, while *M* is the size of the female mosquito population. The mathematical formulation of the SIRS dynamics of transmission is given by the system of equations (1).

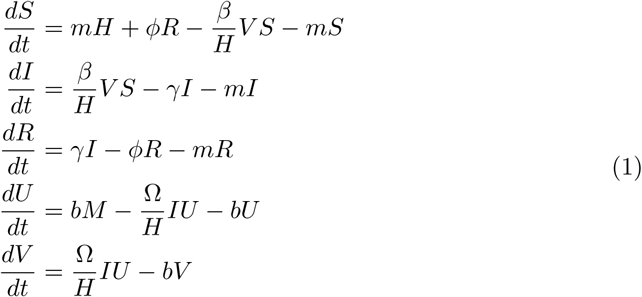

In addition to vector and human basic demographic and epidemiological parameters, it is also common to assume seasonal forcing of the vector population, emulating changing conditions from more to less favorable throughout the year, usually attributable to either hot/cold, or humid/dry seasons. The result of that function is then added either to the birth or death rate of the mosquito population, resulting in a deterministic sinusoidal oscillation.

For clarity the seasonality function is not introduced in this first display of the mathematical system (eqs. 1), but is detailed in the following system (2) instead.

#### 2.1.2 SIR-vector models and multiple infections

Given that dengue virus has a finite number of serotypes, it is expected that any one host can only be infected with dengue a few times in a lifetime; therefore, secondary infections can be modeled by explicit compartments for the secondarily or further infected.

In this case, the total number of times a single individual can be reinfected has a hard limit given by the number of infected compartments in the model, which are never revisited. This is straightforwardly modeled by a series of SIR compartments chained together (i.e. an SIR structure followed by another SIR).

The identities of individual serotypes are not explicit in this model, but sequential infections implicitly describe this feature of dengue virus transmission. The case with two consecutive infections is shown in figure 2.

**Figure 2.**
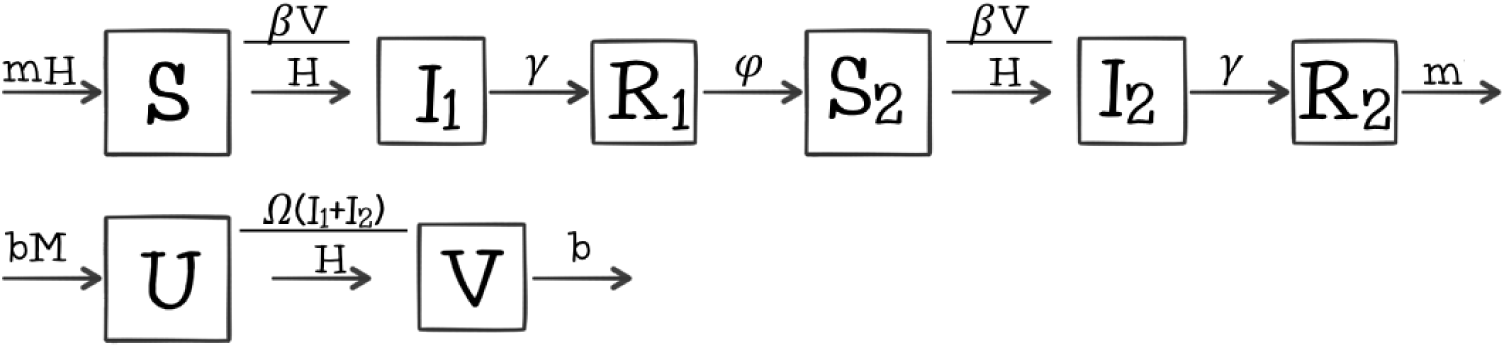
Structure of an SIR plus vector model, with explicit number of possible reinfections (in the particular case illustrated the vector SIRx2).

The compartmental structure of the vector population is unaltered, while the human host population follows a susceptible-infected-recovered path, being immune to reinfection for the time they stay in the first recovered compartment (*R*_1_). After that, loss of immunity takes individuals to a second susceptible state (*S*_2_, unlike the SIRS model where the first state is revisited), where individuals are again susceptible to infection by infected mosquitoes.

In case of infection they move to the secondarily infected compartment (*I*_2_), and after recovery they move to the last compartment (*R*), where they are recovered and can only exit by the ultimate process of death.

The model parameters are the same as the previous model and describe exactly the same processes as before, with the single and only slight exception being that *ϕ* describes a path of waning immunity through different compartments due to the general model structure. The mathematical formulation of the model with two sequential infected compartments is given by the system of equations (2).

As mentioned above, system (2) also has a seasonality term acting on mosquito birth rates. This consists of a time dependent cosine function with argument 2*π* multiplied by time itself plus a phase variable *δ*; this assumes these variables are given in years, resulting in an oscillation period of one year, but can trivially be transformed into months, weeks, or days, for instance, by dividing by the appropriate factor. The cosine multiplies the amplitude scaling parameter *α*; this factor is added to the constant birth rate, resulting in a population whose size oscillates between (1 ± *α*)*M*.

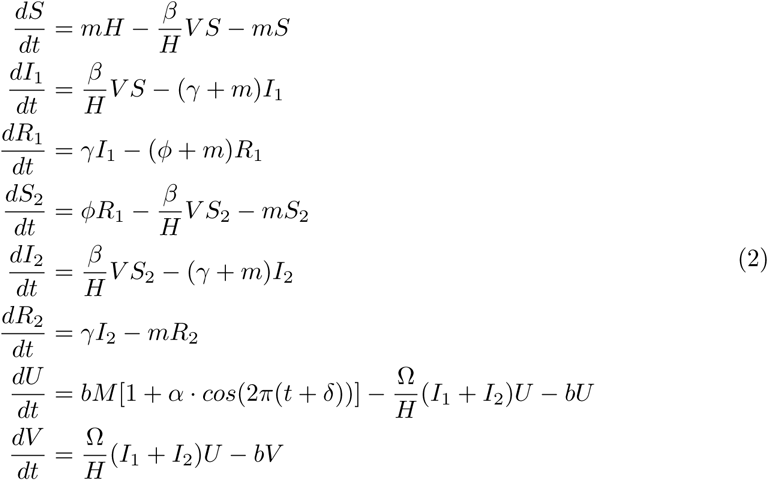

In contrast to the three-compartment SIRS model I refer to this model as the SIRx2 model (given that the human population states are described by a SIR plus SIR, or two times the SIR structure), where once-recovered individuals would again become susceptible to infection. Because *DENV* has 4 human-infecting serotypes infections may further be tertiary or quaternary; unless explicitly stated, I hereafter refer to all infections after the first simply as secondary, as opposed to primary. SIRx3 and SIRx4 models where individuals infected twice or three times, respectively, become once again susceptible can be built by straightforwardly extending the SIRx2; therefore, I do not show specific schemes or systems of equations for those.

In the infinity limit these models become the SIRS model, except human hosts enter an infinite number of new compartments instead of entering the same compartments an infinite number of times, provided of course hosts stay alive long enough. Depending on rates of infection and death, a smaller number of compartments may be enough for a human host to have enough new compartments for many lifetimes of repeated infection, in which case the model will also approach the SIRS model even with a finite number of reinfections.

I refer to this entire class of models as SIRX, which include the shown SIR, SIRS, as well as the SIRx2 (or any other number between two and infinity) models. In any of those cases, however, the identity of multiple serotypes are only implicit in the fact that human hosts can have secondary infections, because there is only one class of infected mosquitoes that transmit to all susceptible humans regardless.

#### 2.1.3 Multiple-serotype infections in the descriptions of dengue virus incidence

A more complete description of multiple-serotype transmission is one that differentiates not only between primary and secondary infections but also disinguishing which serotype causes infection each time. This requires not only a series of compartments, but also parallel paths that describe the order in which the multiple serotypes cause infection. For two serotypes, for instance DENV-1 and DENV-2, it is accepted that a human host could be infected twice, once for each serotype; that is accounted for by the two sequential infected compartments of the SIRx2 model described previously.

Here I wish to account for the order of infection: a human host can either be infected by DENV-1 and then DENV-2, or by DENV-2 and then DENV-1; this creates two alternative paths which are shown in figure 3 – e.g. *I*_12_ denotes individuals first infected with serotype 1 and now infected with serotype 2.

**Figure 3.**
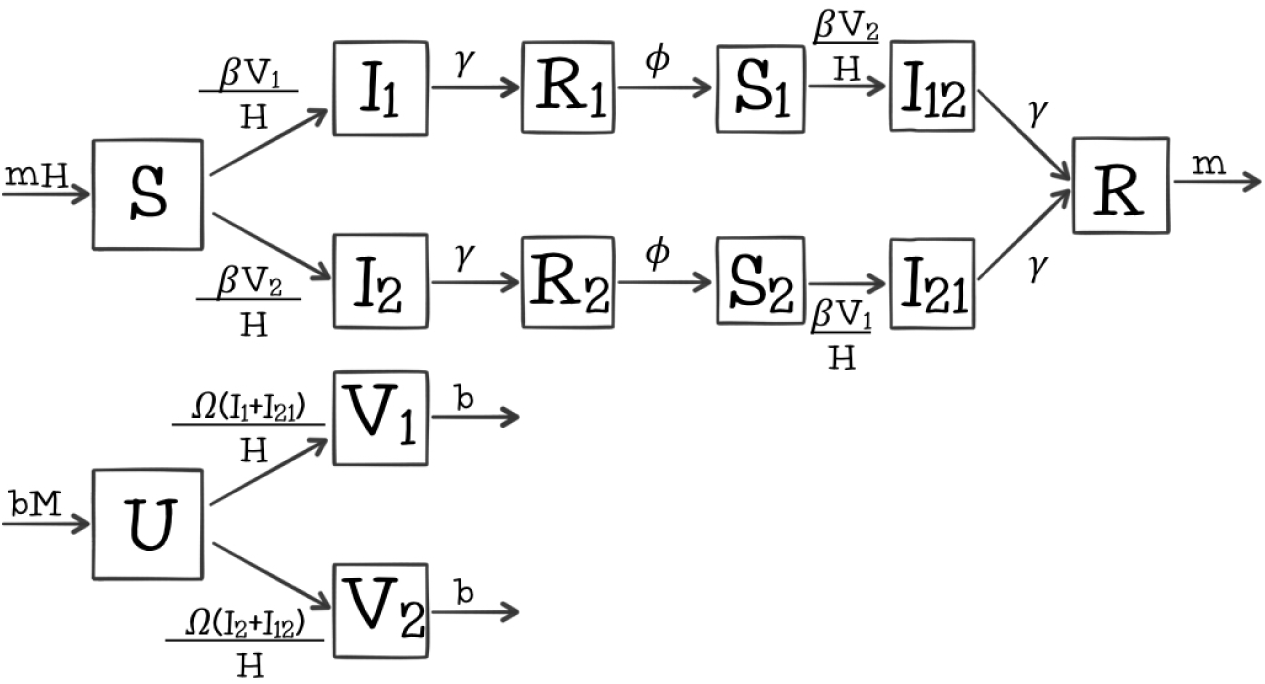
Structure of an explicit two-serotype model and its rate parameters.

Unlike the SIRx2 models the mosquito hosts can either harbour one or the other serotype; therefore, also in contrast to the previous formulations, a secondary human infection depends on transmission from a mosquito with different serotype from that of the primary. The mathematical description of this explicit two-serotype model is given by the system of equations (3).

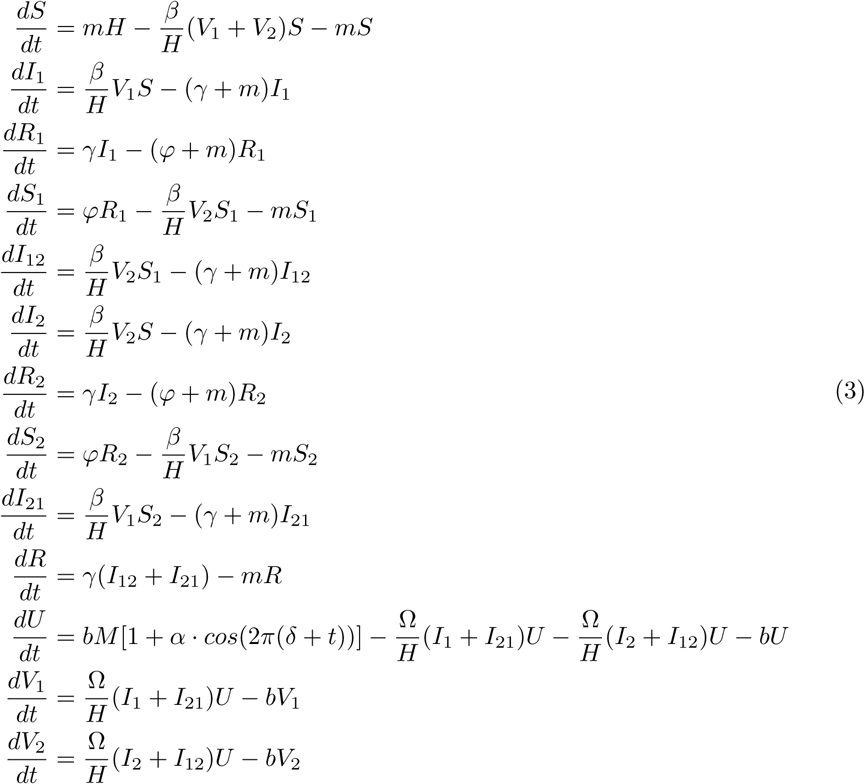

Although the model described by the system of equations (3) does contain (more than a couple) SIR-like components, and could possibly be seen as a parallelization of the SIRx2 model, the level of complexity arising from further introducing *DENV*-specific features is considerably higher. It may be more useful to look at it as two interacting epidemics (Rohani et al. 2003), since it is unlikely that either the compound output or the individual serotype dynamics can be predicted to trivially conform to that of its more well known building blocks.

Regarding the basic reproductive number, *R*_0_, because it is calculated with regard to a fully susceptible population, the secondary compartments do not cause the models to differ in that matter, and the quantity is given by 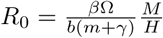. If such differences are actually verified in the output of the model, it would be due to the structure of the immunological states believed to represent a population at risk of a multi-serotype dengue epidemic, since all other processes are present in the previous models as well.

On top of the structuring of basic epidemiological and demographic processes (Keeling & Rohani 2011), additional complexity may appear by introducing asymmetries between serotypes (e.g. serotype 1 more infective than serotype 2, or causing infection for a longer period of infection) or order-dependent rates of infection (e.g. secondary infections more or less likely than primary), which are plausible for various biological or medical reasons, and are likely in comparison to the narrow null hypothesis of perfect symmetry.

Other common extensions that are known to be present to some degree, unlike more complex hypothesized immunological or epidemiological effects, include exposed compartments (describing individuals that harbor the pathogen but cannot transmit it yet), spatial structure of transmission, heterogeneity in contact rates, susceptibility to infection, or in infectivity, gamma-shaped (as opposed to exponential) host survival, and many others. Nevertheless, none of these are included in the models used here, I discuss some of these later in the text, although it may become clear by the section on inference which are the difficulties of including too many parameters however simple and concrete the processes may be.

#### 2.1.4 Implementation

The models were initially implemented as continuous ordinary differential equations (ODEs), solved by numerical methods to approximating the deterministic solutions of the system; commonly available as ODE solver functions in multiple programming languages such as Matlab, Python, and R languages, which were used at different times with no particular preference for either.

### 2.2 Individual-based models

Discrete, stochastic, individual-based versions of the models described above were also implemented. Besides the importance of the randomness of the events in the epidemiological model, it was important to be able to simulate not only the epidemiological outputs, but also the evolution of viral sequences. Because there is no direct continuous approximation neither to the appearance of a random mutation, nor to a genetic sequence of nucleotides, the most straightforward way of simulating evolution is to explicitly attribute viral sequences to infected individuals, and allowing them to randomly acquire new mutations as they get transmitted.

Implementations were done in both C++ programming language by using a previous implementation (Gordo et al. 2009; Gordo & Campos 2007), and later by adapting the algorithm to the Python programming language to take advantage of the random number implementations in the latter.

In brief, an “Individual” class was created to have a “sequences” attribute (which was empty if the individual was not infected), and each human or mosquito host was an instance of that class. At each time step (Δ*t*, which multiplies all probabilities hereafter mentioned), the number of new infections was drawn from a random binomial distribution, since the maximum number of infections is bounded by the total number of susceptible individuals; the probability parameters were equal to the force of infection (e.g. *λ* = *βV*_1_*/H* for infections caused by mosquitos infected with serotype 1) and number of trials equal to the susceptible population (e.g. the susceptible to all *S*, or to type 1 *S*_2_, equivalently.) Because the infectivity of all individuals is assumed to be the same, the sequence infecting each new host is randomly drawn from the pool of all existing sequences from the previous time step. After a successful infection of a new individual, new mutations have the opportunity to arise with probability determined by a per-genome mutation rate – mutations are assumed to follow an infinite alleles and sites model, so every new mutation is one that was not previously present in the population. Mutations do not affect any of the model parameters, so evolution is completely neutral. The number of sequences inside a single infected host can be greater than one, in which case this within-host population undergoes a Wright-Fisher sampling step (Wakeley 2009).

The number of new births of the mosquito population, apart from the sinusoidal additional factor, is drawn from a poisson distribution with mean *µ*_*vec*_ = *b*; deaths are drawn from a binomial distribution with probability parameter also *µ*_*vec*_ = *b*, so the total population is expected to fluctuate around the initial value. The human population is assumed to be strictly constant; that is enforced by the number of births being exactly equal to the number of stochastic deaths – this condition can be easily relaxed, however.

Otherwise, as a general rule, the number of events at each time step was drawn from a random binomial (and when applicable, multinomial) distribution where the number of trials was given by the number of individuals in the compartment, and the parameter for probability of success in each trial was given by the rates in the model (e.g. the number of individuals recovered from an infected compartment *I*_1_ is given by a random binomial distribution draw with parameters *I*_1_ and *γ*). If competing processes were present, a multinomial was used instead. If the number of events was not bounded, for instance by the size of the compartment, a poisson distribution was used instead (e.g. the number of mosquito births, or number of new mutations).

The output of this implementation is both a stochastic time series of susceptibles, infected, recovered, and incidences (i.e. the randomly drawn number of new infections recorded at each step in the human and vector populations), and the pool of all extant pathogen sequences for each serotype at all or selected time points (for convenience, split into mosquito and human harbored sequences).

### 2.3 Epidemiological surveillance data (and pseudodata)

In countries like Brazil, where notification of dengue is compulsory to doctors, the records commonly consist of periodically reported new cases into the Information System for Incident Notification [Sistema de Informação de Agravos de Notificação] (SINAN). As with many other common endemic diseases, laboratory confirmation is not routine, so diagnostic relies mainly on clinical criteria; neither the serotype causing the infection is normally recorded, nor if the infection is primary or not. Therefore, typical time series do not distinguish between serotypes or sequential infections; what would be available would be a series of equally spaced, discrete values representing the number of cases of “dengue fever” generically defined reported every month or week (SINAN).

The time series data set used here is from the Brazilian SINAN, with the absolute number of new weekly cases of dengue from the year 2009 until 2013 in the city of Rio de Janeiro, when three large incidence peaks are observed. More specific diagnostics data have also been occasionally produced in the form of serological surveys, although these were obtained for specific studies and small cohorts. Because this kind of data is sparse and difficult to access, I do not use any such data, but merely note that it exists for the disease and location I am (mainly) concerned with, even if in a fragmented way.

Relatively recently, routine surveillance of dengue started to include genetic data of the virus. It is now routine activity to isolate samples from patients and obtain nucleotide sequence from the virus in the isolate; an immediate result is the identification of serotypes, and possibly of more specific variants such as genotype (a finer grained distinction within each serotype) as well as the relationship to strains previously found elsewhere (dos Santos et al. 2002). This data set therefore consists of somewhat sparsely sampled viral isolates sequenced along several years – the total number is in the order of tens for the city of Rio de Janeiro – and is to a great extent available as part of published studies (Araújo et al. 2009; de Bruycker-Nogueira et al. 2015; Castro et al. 2013, 2012; Miagostovich et al. 2003, 2006; DeSimone et al. 2004), as well as in public databases for genetic sequences such as GenBank (Benson et al. 2015).

The individual-based models described in the previous subsection were designed in a way that could directly reproduce the form observed in the real data available. Unless there is specific interest in greater details, whenever a model is simulated I try to store the output in a format that mimics the amount, type and level of aggregation, period and interval of collection, and any other feature pertinent to a specific data set. When used in the same way as the real data (e.g. for parameter inference), I call a data set of this sort synthetic, or pseudo-data.

Generally summarizing the two types of pseudo data sets used here, the time series are weekly records of all new cases (the weekly sum of daily-generated incidences), and the second type are genetic sequences. The genetic equivalent of the all-inclusive time series data would be a sequence (or group of sequences) for every newly infected individual time-stamped with the week (or any other time step) when it appeared. That is impractical even for simulated data set, and for a series of reasons it is essentially impossible in real epidemics.

Instead, the pseudo-sequence data is a sample from different times, where the number of sequences from each time point is proportional to the number of cases then, i.e. it is a sample of sequences over one long period, weighted by epidemic size at each of multiple small intervals. A total number of 100 sequences per serotype over the course of a few years was assumed to be a sufficient number, comparable to that of previous data sets used for similar purposes (Rasmussen et al. 2014b), and considered feasible in a city with a population on the order of 10 million, and outbreaks on the order of a few tens of thousands (SINAN). It is also comparable (though larger) in size to data sets collected for other purposes in the city of Rio de Janeiro (DeSimone et al. 2004) When needed, the specific pseudo and real data sets are detailed at the pertinent results sections.

The objective is to obtain pseudo data sets suitable for inference purposes. It is difficult to establish beforehand what the most informative sampling scheme would be (Frost et al. 2015); that is a question on its own right that is not explored here.

### 2.4 Bayesian inference from time series

Analytical solutions for the systems adopted in this chapter are not available; therefore, a numerical approximation to the continuous solution was obtained through an ODE solver whenever needed.

Because the number of new cases in any given week is an integer number I chose to use a poisson distribution: the likelihood of the observed value is computed using the sum of all possible human infected states as the poisson parameter (as modeled by compartments, i.e. the total number of dengue cases of any kind in the model output), and the total likelihood is therefore the product of that over all time points in the series – or, more conveniently, the sum of their logarithms.

A binomial distribution could as well be used, in which case its probability parameter could be given by the forces of infection and the number of trials would be given by the susceptible populations at risk. While that would be possible, and possibly mimic more accurately our simulation model and the bounds in the maximum number of infections, it is more cumbersome to add the parameters coming from the different compartments and, more importantly, the poisson distribution expects greater or equal variance when compared to the binomial, and therefore can accommodate any overdispersion in the data.

The bayesian Markov Chain Monte Carlo (*MCMC*) inference algorithm was implemented in Python language using the PyMC module (Patil et al. 2010). Unless detailed otherwise, as a norm gamma-shaped priors were used for parameters that have independently estimated or commonly accepted values, and priors uniform over a wide range were used otherwise. As technical criteria for quality of the inference, the following criteria were used (Gelman et al. 2013, chap. 11): Markov chains were run until the likelihood and posterior traces converged to a maximum and attained stationarity with that regard; for all results shown, replicates of the chains were run to assure mixing (unless otherwise specified, model fit and posterios are computed and shown for individual chains only); initial sampling corresponding to one tenth of the total number of iterations was discarded as warm-up or burn-in period. Correlation between parameters were computed at the end of the chain for at least one replicate when parameters seemed systematically biased.

Most critical was the time limit of around one month that was imposed for practical reasons; that allowed chains as long as a few million iterations on a dedicated high-capacity computer. Most implementations did not seem to have issues with the likelihood converging with a few hundred thousand iterations; nevertheless, more subtle issues were observed in some cases and are discussed in the results and discussion sections.

I applied this estimation method to both simulated data sets, as well as to a time series of dengue incidence from the city of Rio de Janeiro running from 2009 to 2013 (SINAN).

The estimation method used does not therefore account for noise in the system state. Stochastic solutions can be used instead; however, doing so is not as simple as replacing a deterministic solution by a single stochastic one (which may be only slightly slower to obtain), but requires a considerably more sophisticated and a computationally much more intensive method method to estimate the likelihood Andrieu & Doucet (2010); Ionides et al. (2006), with possibly a couple of thousand simulations at each iteration of the Markov Chain. I discuss these so called Sequential Monte Carlo or particle filters as perspectives in the end of this chapter.

### 2.5 Population genetics and phylodynamic inference

Although the individual-based model can produce a simulated data set that mimics a set of viral sequences sampled at arbitrary time points along time, unlike with the time series data there is no simple way of calculating the likelihood of the model parameters given that kind of data.

The conceptually most straightforward method is probably the following: make up metrics that are assumed to be representative of the data; simulate the model; compute the same metrics for a large number of model outputs; and try to find the model parameters that best approximate the real data. In spite of the gross oversimplification of this description, this is the basis of Approximate Bayesian Computation (or *ABC*) methods.

An alternative framework to compute the likelihood of a model given sequence data – and arguably a more elegant one, at least in the sense that it is based on a full likelihood expression – relies on the bifurcating properties of the trees that connect related sequences, or conversely (with time flowing backwards) the coalescence of a set of related samples into a common ancestor. The latter gives the name to the coalescent theory, or simply, *the coalescent*, as described by Kingman (1982). Since then the problem of estimating a Wright-Fisher (or Moran) population size from sequence data using the coalescent has been extended to varying environments (Griffiths & Tavaré 1994), to implicitly defined population functions (Frost & Volz 2010), and more generally to structured populations (Volz 2012).

The Beast 2 software (Bayesian Evolutionary Analysis by Sampling Trees) contains many basic as well as advanced implementations of coalescent-based *MCMC* inference (Bouckaert et al. 2014); it is written in the Java programming language, and uses an XML file to input the actual sequence data and the specifications to the multiple classes involved in the calculations. A phylodynamics package is also available, which among many things includes implementations of the SIR model (Kühnert). Other software for that purpose is also available, notably as packages for the R programming language (Paradis; Volz et al.). Beast 2 was chosen due to the existence of a general community of users, its openness and extensibility, apparent propensity to phylodynamics implementations, and especially the helpfulness of some of its earliest developers.

Nevertheless, there were no tools built in to the core software, nor any extension packages (including the phylodynamics package developed by Kühnert) that were suitable for direct application to the inference using the models pertinent to dengue virus transmission as describe above. For one, a structured implementation is needed – in the simplest case, viral sequences may be either in the human or in the mosquito host, and intuitively coalescence can only happen if they are inside the same hosts (unfortunately, further details about structure in the coalescent for this kind of model are way out of scope, but see Volz (2012) for a complete description of how structure is incorporated in the context of disease transmission models). The implementation was greatly facilitated by code shared from a previous implementation from Rasmussen et al. (2014b), which consisted of an epidemiological model Java class and a structured coalescent likelihood computation class; these extensions could almost directly be applied to the SIR-vector model, and could straightforwardly be used for the SIRS model as well, and more or less easily adapted to all SIRX models.

The implementation of the two-serotype models, however, required additional tinkering. Because *DENV* serotypes are only 60% similar by sequences, they are usually treated separately (de Bruycker-Nogueira et al. 2015; Castro et al. 2013, 2012). Therefore, they are not expected to find a common ancestor in the recent period of a few years of dengue transmission, in which case two separate trees are needed. The two-serotype epidemiological model produces the poulation dynamics function for both serotypes, so a single Java class was implemented with this model; however, the likelihoods have to be calculated for each tree, so two separate classes were implemented that used separate trees and substitution model parameters, but fetched (different, serotype-specific parts) of the same population model output. The combination of the two tree likelihoods was then the global likelihood of the one-epidemiology, two-tree model, and from then on could be used by the *MCMC* algorithm in Beast 2 with no additional tinkering needed to perform inference. The same quality checking criteria as in the time series inference were used; one month of real time amounted to a few tens of millions of iterations in Beast 2 for the most complex models. Additionally, Beast 2 computed the effective number of samples (Gelman et al. 2013, chap. 11) as an additional metric to assess convergence.

## 3. Results

### 3.1 Forward model simulation

#### 3.1.1 Predictions for disease incidence and prevalence

Figure 4 shows a deterministic simulation of the SIR-vector model without, and with seasonal forcing in the birth rates. It is worth noting, that although the amplitude of seasonal forcing is of the magnitude of 10%, oscillations can quite easily be greater than that through resonance of the natural (damped) oscillations and the forcing (Dushoff et al. 2004). The average incidence, however, is roughly the same in the presence or absence of forcing.

**Figure 4.**
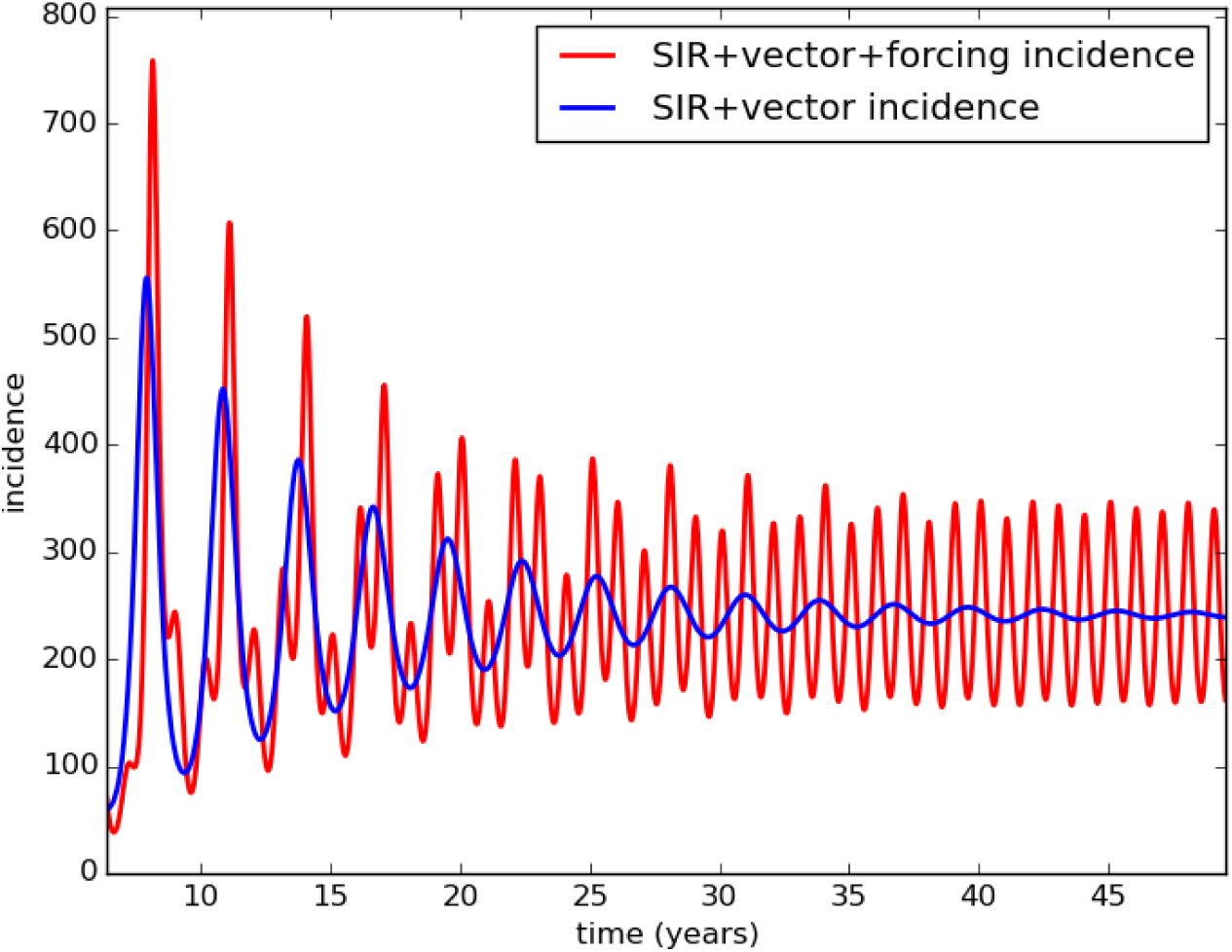
SIR+vector models with (red) and without seasonal forcing (blue) Parameter values in are: Ω = *β* = 1.0, *b* = 0.1, *m* = 3.65 10^−5^, *γ* = 0.14, *α* = 0.1, *H* = 1000000, *M* = 513789.

In the case of waning immunity, secondary infections are also possible, which potentially increases the total incidence at any point in time. Figure 5 shows the total incidence for the vector transmitted SIR, SIRx2, SIRx4, and SIRS models; for a set of parameter values, changing only the transmission rate within a certain range can result in a very similar incidence profile even in a deterministic setting.

**Figure 5.**
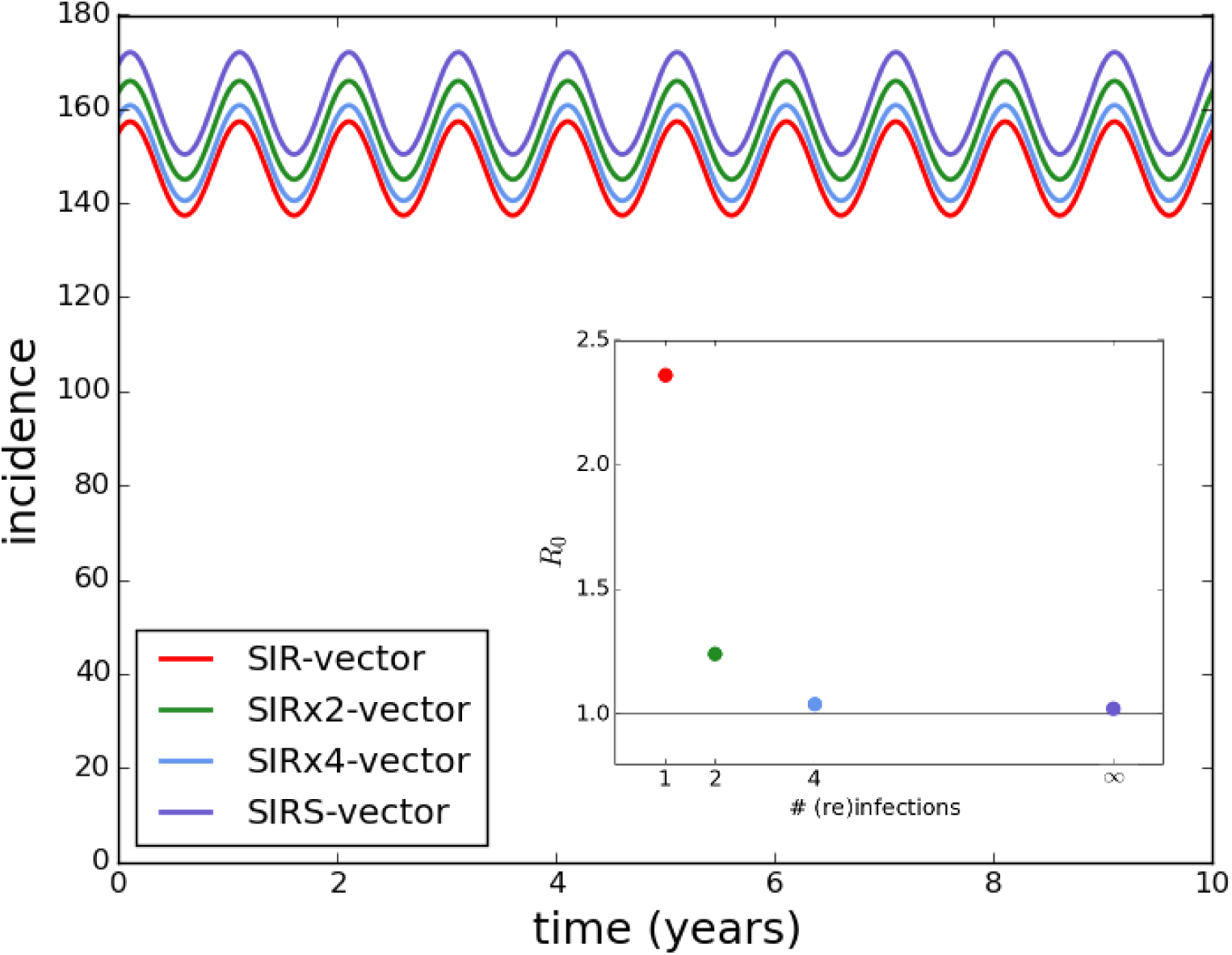
SIR+vector models in some of its variants: standard vector SIR (red), SIRx2 (green), SIRx4 (light blue), and SIRS (purple), showing approximately the same incidence levels at or near near the oscillatory equilibrium. The single changing parameter is transmission intensity for the SIR, SIRx2, SIRx4 and SIRS models, respectively: Ω = *β* = 0.2536; 0.1835; 0.1680; 0.1665. Additional parameter values in are: *b* = 0.1, *m* = 3.65 · 10^−5^, *γ* = 0.14, *ϕ* = 0.00165, *α* = 0.02, *δ* = 0, *H* = 1000000, *M* = 513789.

Although for larger transmission rates the number of susceptibles may dominate and cause the models to output distinguishable time series, it must be acknowledged that for some parameter sets, model structure is not easily identifiable from the incidence pattern. Therefore, not only parameter values, but also model structure, are likely to be critical to describe the transmission of dengue virus, and can have a great impact in the parameters estimated.

The structure of the two-serotype model is not directly comparable to the previous models; nevertheless, the output of the model with the same common set of parameters (except for *β*) is shown in figure 6, except for transmission.

**Figure 6.**
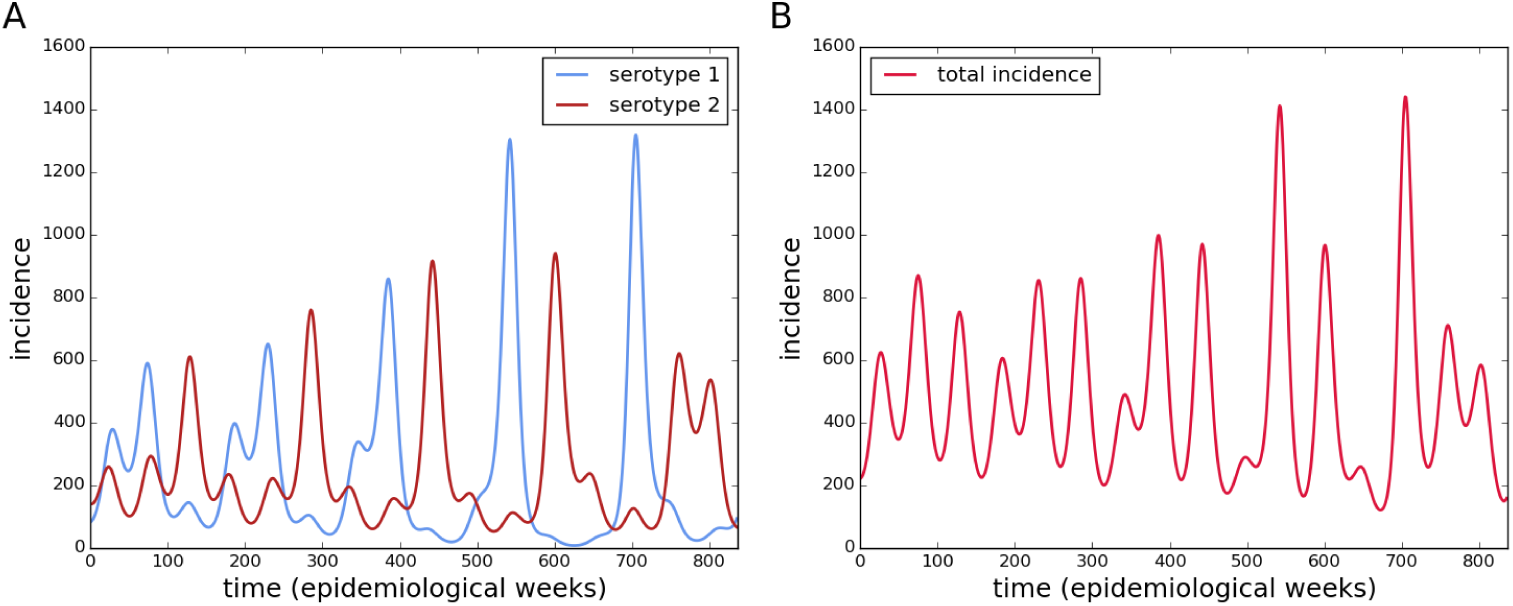
Incidence dynamics for an explicit two-serotype dengue virus model. Shown incidence is that for each serotype separately (a), and the sum of new infections for both serotypes (b). Parameter values are: *b* = 0.1, Ω = 0.7, *β* = 0.7, *m* = 3.65 · 10^−5^, *γ* = 0.14, *φ* = 0.00165, *α* = 0.02, *δ* = 0, *H* = 1000000, *M* = 513789.

Besides the obvious difference that this model has explicit series for both serotypes, the incidence pattern is less regular. A more striking qualitative difference is the fact that the numbers of infected individuals get much closer to zero (at least for individual serotypes); in a deterministic model it means just that, but in a stochastic model that may mean increased probability of extinction.

#### 3.1.2 Discrete time and individual-based implementations of vector transmission of DENV

Discrete time and stochasticity in the system state transitions can significantly affect the oscillations in resonating systems (Dushoff et al. 2004). This is illustrated by the incidence outputs of both the vector SIR, and more extremely the vector SIRS models, in figure 7.

**Figure 7.**
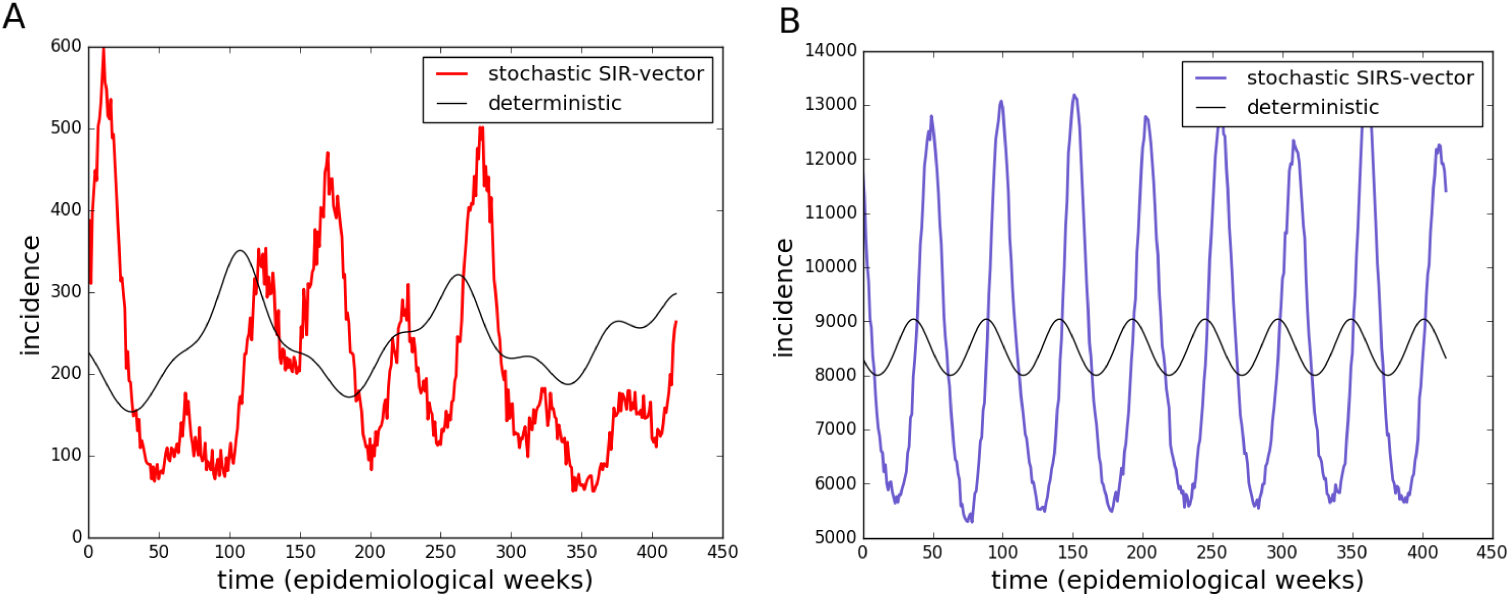
Stochastic output of vector SIR model (a), and of SIRS model (b). Gray lines show solutions to the deterministic system. Parameter values are: *b* = 0.1, *m* = 3.65 10^−5^, *γ* = 0.14, *ϕ* = 0.00165, *α* = 0.02, *δ* = 0, *H* = 1000000, *M* = 513789., and Ω = *β* = 0.7 for the SIR, and Ω = *β* = 0.47, for the SIRS.

The probability of extinction of the pathogen is generally low in those cases, although it is clear that numbers are much lower for the case without reinfection (vector SIR), such that if extinction doesn’t commonly occur, the trajectories are nevertheless much more prone to the general process noise. Similarly, the intermediate models with two and four infections are prone to effects of discretization and stochasticity. Figure 8 shows the results for the remaining SIRX models.

**Figure 8.**
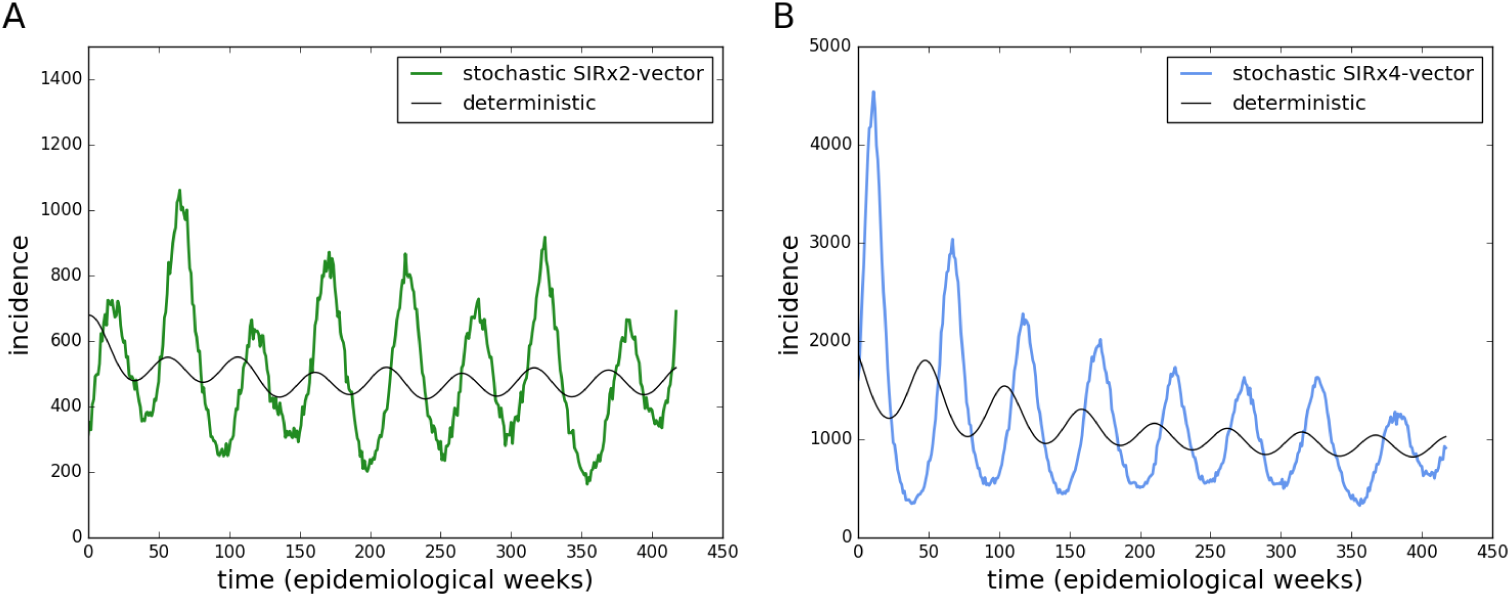
Stochastic output of vector SIRx2 model (a), and of SIRx4 model (b). Gray lines show solutions to the deterministic system. Parameter values are: *b* = 0.1, Ω = *β* = 0.7, *m* = 3.65 *·* 10^−5^, *γ* = 0.14, *ϕ* = 0.00165, *α* = 0.02, *δ* = 0, *H* = 1000000, *M* = 513789.

For the chosen parameters, extinction is more likely in the two-serotype model Figure 9 shows the results for this model.

**Figure 9.**
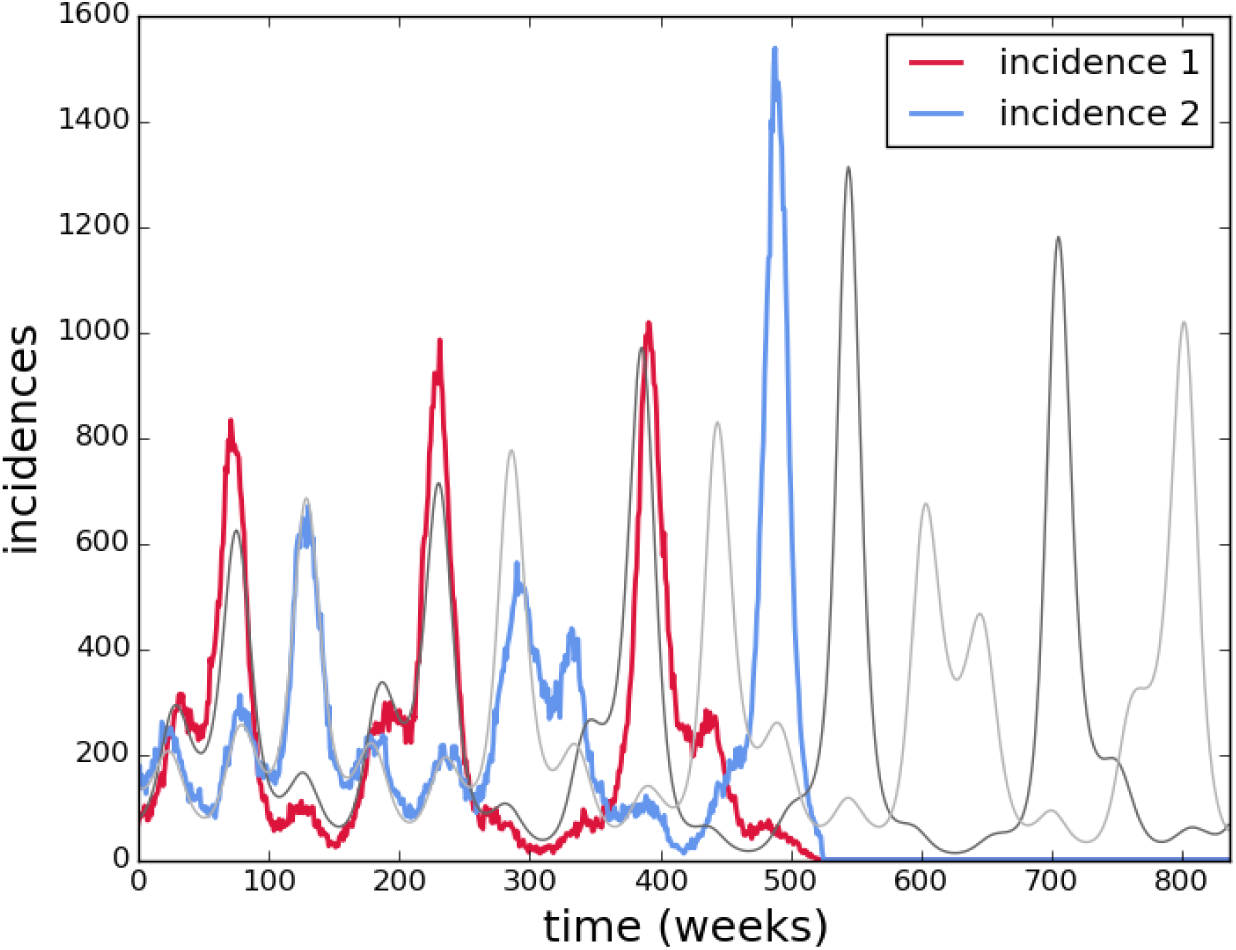
Stochastic output of two-sreotype model. Gray lines show solutions to the deterministic system. Parameter values are the same as before.

#### 3.1.3 Summary of forward modeling predictions

Models that are structured differently but with the same parameters, and therefore same basic reproductive number, will have different outputs and observed incidence and prevalence levels. Conversely, qualitative aspects of the observed disease data can be reproduced by different combinations of model structure and parameters. It is not at all trivial to determine what aspects of the model are known, or best approximate reality, which can be treated as nuisances, and which of them are robust to structure or parametrization changes.

In the case of genetic diversity, it is even harder to distinguish clear relationships between the real data and the model predictions. Summaries are useful to make broad assessments about the data, but are usually not suitable for finer grained comparisons, nor model testing. A quantitative comparison is therefore needed to assess which model best represents reality, and what parameter values explain the processes of disease transmission.

The forward simulation approach is quite convenient when good estimates are available for all or most parameters, and there is good confidence in the model structure, or the outputs are robust or easy to evaluate for unknown parameters. Even for a reasonably large amount of synthetic (or pseudo-) data it is feasible to perform computation of genetic summaries in a reasonably short time without much algorithm optimization effort.

For complex, non-linear models it may, however, be difficult to thoroughly perform sensitivity analyses for more than a couple parameters and identify disease dynamics compatible with real data. For genetic diversity computations that rely on individual-based simulations it is even more costly and time-consuming to adopt the forward approach.

### 3.2 Inference: quantitative hypothesis testing

#### 3.2.1 Time series-based inference

The *MCMC* algorithm produces samples of the joint posterior, which consist of a series of parameter sets accepted by the method, the fit can be empirically computed by simulating the model for a representative subset of these samples, and computing the mean or median values of the model output. Credibility intervals can be similarly computed by taking the score at some percentile (e.g. a 95% CI results if 2.5% and 97.5% are chosen). We present the mean computed as described above and call it the “fit” hereafter, together with the 95% credibility intervals, unless otherwise stated.

For inference purposes, the ratio of mosquito population *M* to human hosts *H* is the estimated parameter, denoted hereafter as *M*_*ratio*_. The mean time of immunity before becoming susceptible, that is 1*/φ* or 1*/ϕ* are the parameters actually estimated, and are denoted just as such, the inverse of the original parameter.

The easiest data set the inference method can fit is a deterministic simulation with some noise added afterwards to the continuous solution. This best case but unlikely scenario serves as proof of concept that there would be enough information in a data set with this format. Figure 10, panel A shows the fit of the two-serotype model to pseudodata produced by a deterministic simulation of 209 weeks (approximately four years) with poisson noise added to each of the 209 time points, i.e. the model used for estimation is the same as that used to produce the simulated data.

**Figure 10.**
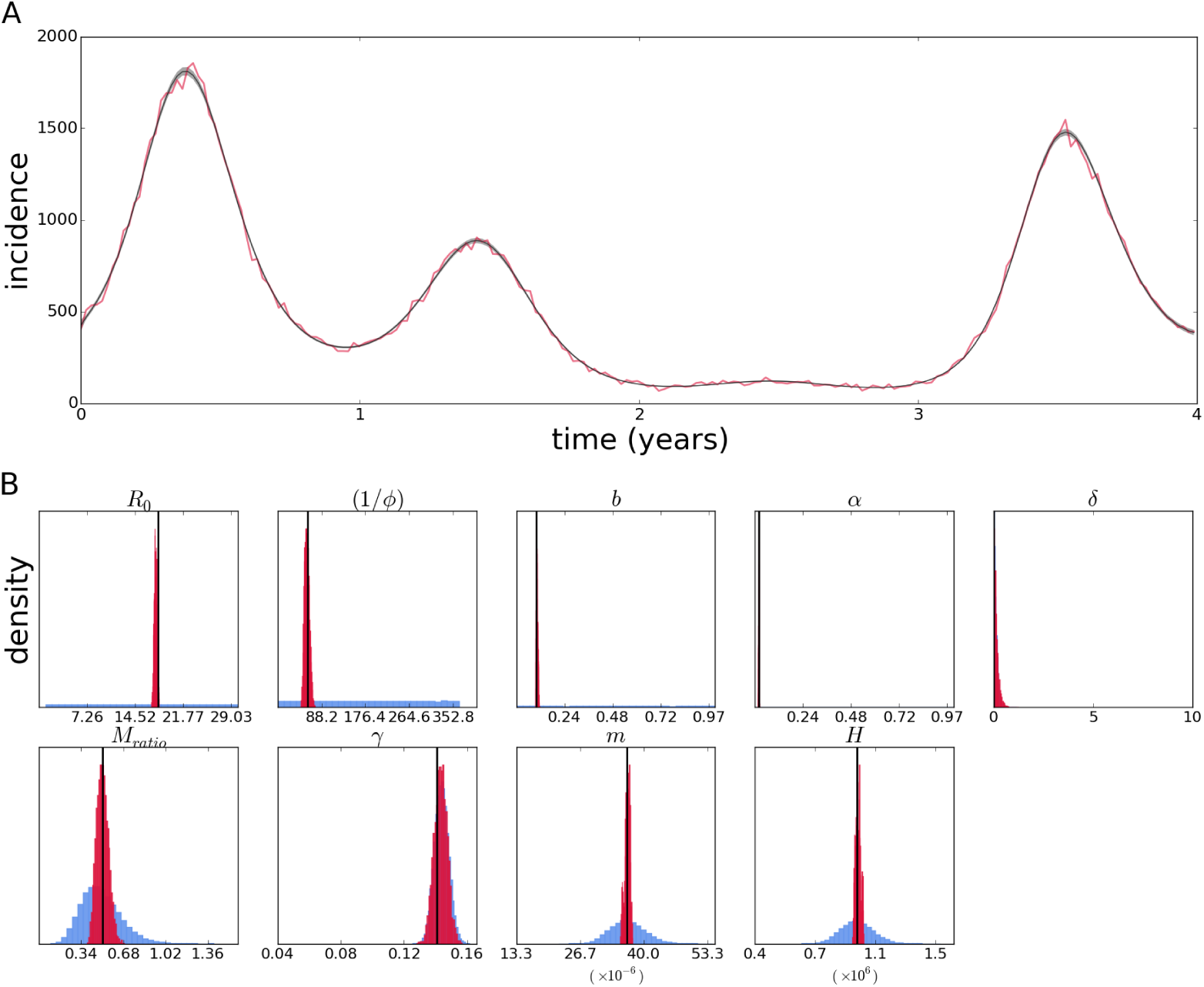
Fit of the continuous two-serotype model with poisson likelihood to data simulated from the continuous model with poisson noise added afterwards (A). Posterior distributions (crimson) of epidemiological parameters (B) – prior distributions are shown in light blue (possibly not visible if the densities are very low compared to the posterior density). True values, i.e. parameter values used to simulate the data set, are shown as vertical black lines.

The fit shown in is extremely good, considering the pseudodata is the output of a highly nonlinear model, and that only an aggregated and partial observation of the system is used (i.e. the sum of the changes in the infected compartment). Therefore, while this is the best case scenario, it is not a given that it would be possible to adequately fit the model, and furthermore estimate the parameters accurately – the parameters could be structurally unidentifiable, or the amount or type of data could not allow accurate estimation of the parameters. The estimates of the model parameters are shown in figure 10, panel B, which displays the prior and posterior distributions. Most estimates are quite precise even in the absence of informative priors.

The case of the exact same estimation method applied to a data set produced by the stochastic individual-based model simulation instead is shown in figure 11. While the fit to the data is still quite good, the effect of stochasticity on the posterior estimates can be observed as pronounced biases in some posterior distributions.

**Figure 11.**
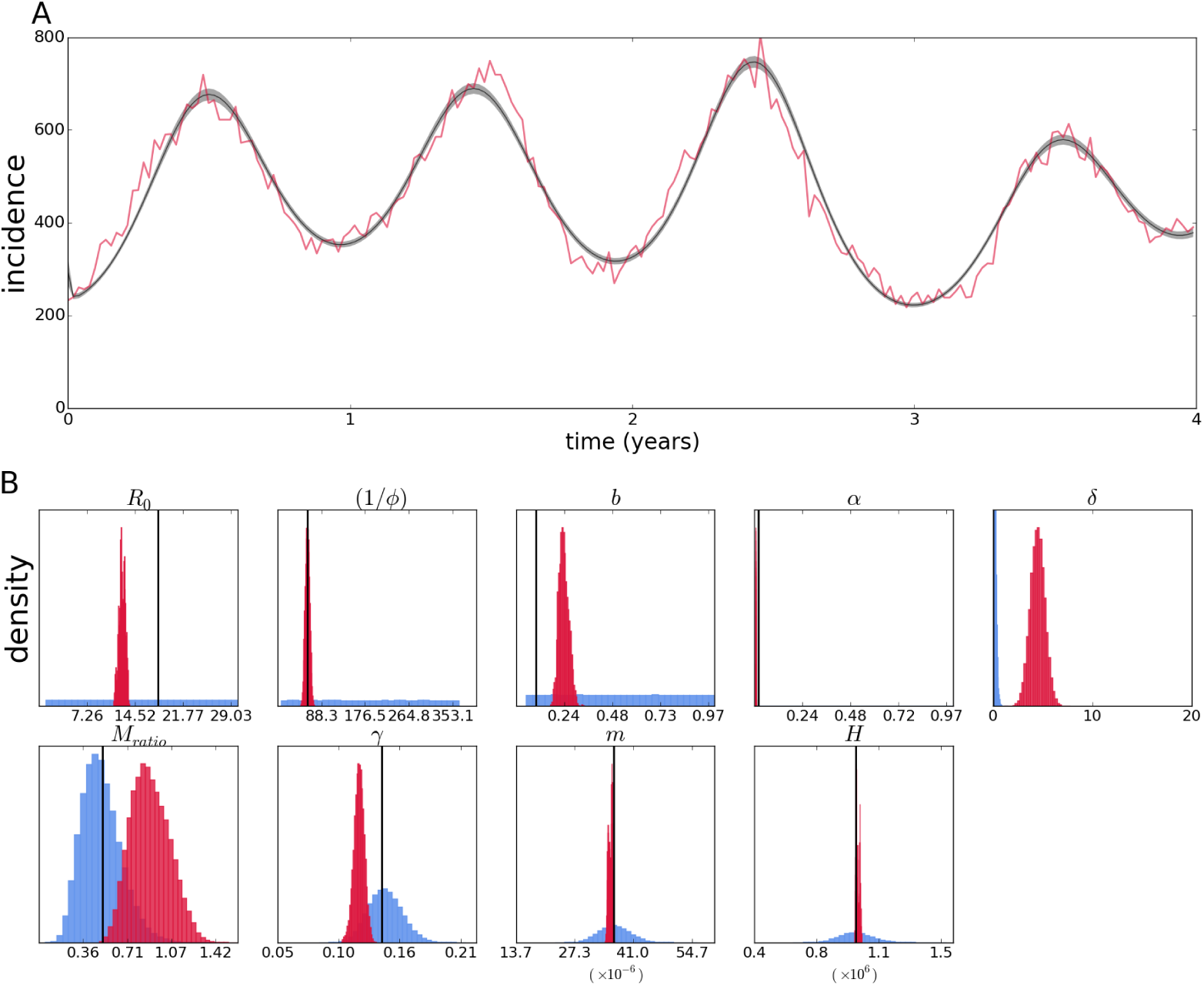
Fit of the two-seroytpe continuous model with poisson likelihood to data simulated from its stochastic version (A). Posterior distributions (crimson color) and priors (light blue) of epidemiological parameters and initial conditions (B).

Unlike the previous case, gamma priors were used for the initial population states, and they are nevertheless not as well estimated as before, but contain the true values (not shown). Particularly important is the estimate for *R*_0_, which is significantly lower than the true value and the estimate of the temporary cross protection time (i.e. strain-transcending immunity – the inverse of the rate of immunity waning *φ* – only present in models with secondary infections), which is precisely estimated.

The observed biases are similar to those observed with a simpler vector SIR model (see appendix,), and the fact that these are not observed with the continuous simulation suggests that model complexity or data structure are not the main factors.

Indeed, fixing some parameters and/or initial conditions can improve estimation of the remaining parameters. Appendix A shows that fixing all other epidemiological parameters except for *R*_0_ improves it estimate and fixing initial conditions improves it further (figure A.2) However, it is not necessarily true that the more parameters fixed, the better the estimates, as leaving a few of the epidemiological parameters results in the best estimate of *R*_0_ when compared to the above (figure A.3).

The *MCMC* algorithm allows empirical computation of the correlation between the parameters, since it relies on repeatedly sampling the posterior distribution of parameters. The biases could therefore be at least in part attributed to correlation between the estimates (not shown, but see discussion section); however, the problem seems to go beyond lack of identifiability of specific parameter combinations, since allowing some parameters to accommodate uncertainty can improve inference results. I further discuss parameter correlation, fixing parameters, uncertainty in estimates, and possible methods to get around these issues in the next section.

Because incidence data most likely does not differentiate between infecting serotypes, a time series gives no information about the number of circulating serotypes, and therefore it is not possible to decide on single or multi-serotype models based on that alone. For inference purposes a number of circulating serotypes must be assumed when a transmission model is used.

The previous results assume the correct model structure (although not the correct error model) is known; with real data other factors such as model misspecification can introduce further sources of errors. Appendix A shows that using the vector SIRx2 model to estimate parameters from a time series simulated from the two-serotype model can throw estimates off (particularly that of *R*_0_); using the same model on data simulated from the vector SIR model can have equally problematic results. That highlights the importance of testing different alternative models for the same data set, when it is not possible to favor any specific model based on the data alone.

It is in principle possible to distinguish the infecting serotype, and even the order of infection, for time series data; however, that would require elaborated tests and/or record keeping for current and previous infections for every single recorded case. Sequence based inference relies on a sample of cases, not all possible records, and differentiates infecting serotype by default, which can get around some of these issues.

#### 3.2.2 Genealogy-based inference

The aim of this section is to show results of methods comparable to that of time series-based inference. In Rio de Janeiro, the available dengue virus sequence data seems to be a result of concentrated efforts to obtain data representative of each single epidemic period, not an overall data set with particular features. To my knowledge, there is no specific goal for the entirety of the available data as to the total size, sampling interval, or representativity of both epidemic and inter-epidemic periods, for instance. Therefore, the simulated data set used for inference here is created to mimic more of an ideal yet feasible data set to collect. In practice that was done by deciding on the total number of sequences desired and sampling with a constant probability that would yield that expected value – given that binomial sampling was done every week, periods with greater number of cases would be more represented in the data.

For a constant probability to be specified, the total number of cases in the entire interval should be known before sampling, which cannot be the case in the real world; even if that was known a probability of sampling a sequence cannot be directly decided on by researchers. Nevertheless, the computational sampling scheme mimics the real world in that the greater the number of infected individuals, the more report to hospitals and health centers, and the probability of obtaining consent to get biological samples and sequencing them can be set by an arbitrary rate (e.g. a goal to obtain a sample from one out of 100 patients) that would yield a total number of samples over a usual epidemic.

For the vector SIR model, a pseudodata set was sampled from the last four years of a sixteen year run resulting in 106 simulated sequences with random mutations. The results from estimation based on that data are shown in figure 12. The posterior contains the true values of the parameters, although some are slightly biased, not centered around the true value.

**Figure 12.**
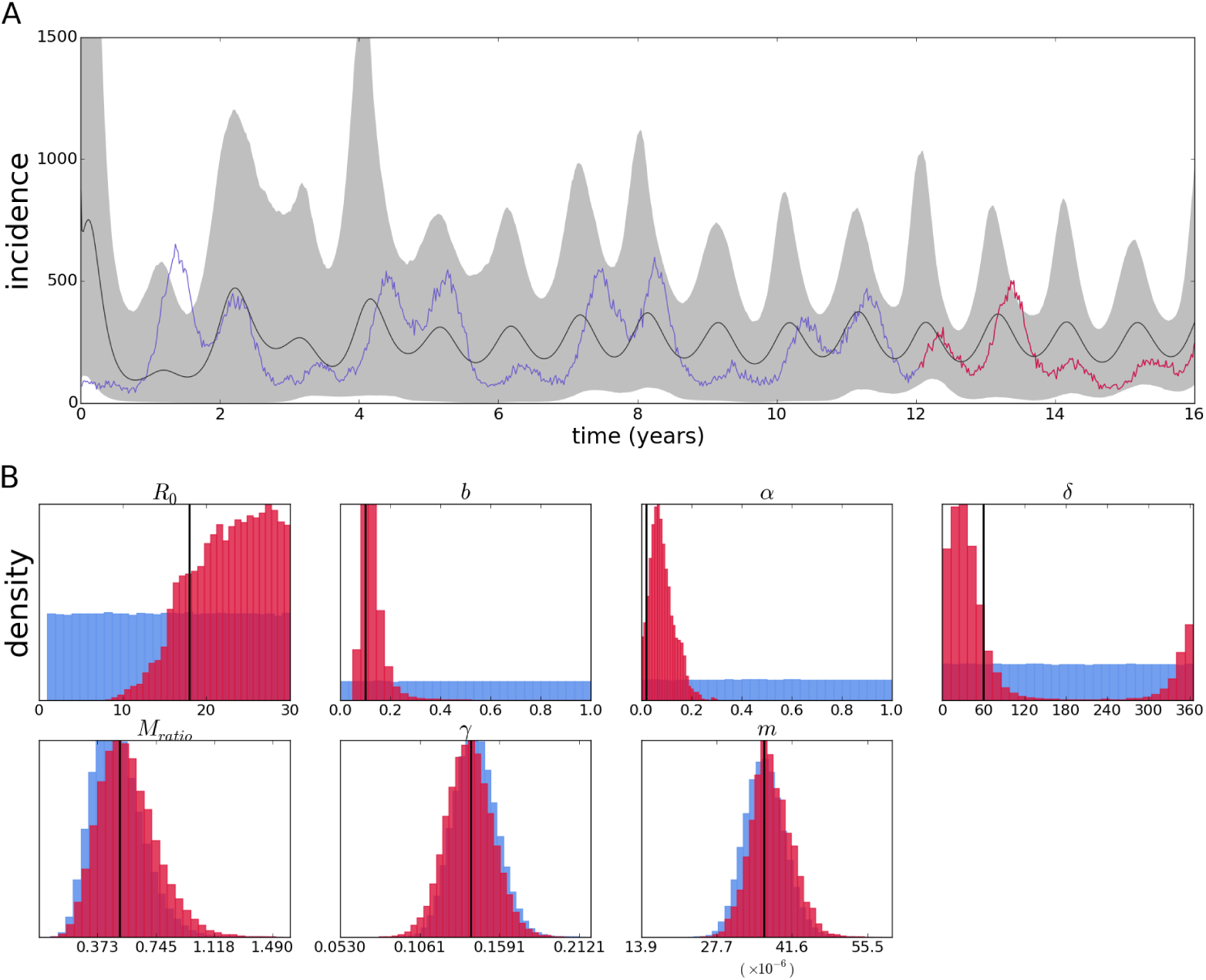
Reconstruction of the epidemic with a vector SIR model from sequences simulated by its corresponding individual-based model(A). Red portion of the time series denotes period from which samples were taken. Posterior distributions (crimson color) and priors (light blue) of epidemiological parameters (B).

Again an expected value and confidence intervals are computed from the sets of parameters in each step of the Markov Chain; however, figure 12A does not show a fit, since the time series data is not used for estimation, but rather a reconstruction of the incidence patterns. Nevertheless, this reconstruction may accurately depict the mean incidence if the sequence data is informative enough, and here it indeed reproduces oscillations on the same order as those in the data. The phase of the oscillations are not perfectly synchronized, and the amplitude is more regular than the actual incidence (although this is expected due to the stochastic data as opposed to the deterministic nature of the inference method; this is further discussed in the next section).

For the two-serotype model, the pseudo-sequences used here for inference were produced by the same run of the individual-based model as the time series pseudodata (in the previous subsection) – as would be the case with a real epidemic, where all possible new cases are recorded as incidence, and some of them are sampled for sequencing. The pseudo-sequence data is sampled from the same 209 weeks for which pseudo-incidence time series was recorded; the total simulation length was 16 years (therefore, the pseudo-time series in the previous subsection actually consists of the last 4 years of this longer run. Only the sequence data is used to produced the results that follow.

Inference using 119 type 1 sequences and 104 type 2 sequences sampled according to the above scheme from a population of simulated individuals is shown in figure 13.

**Figure 13.**
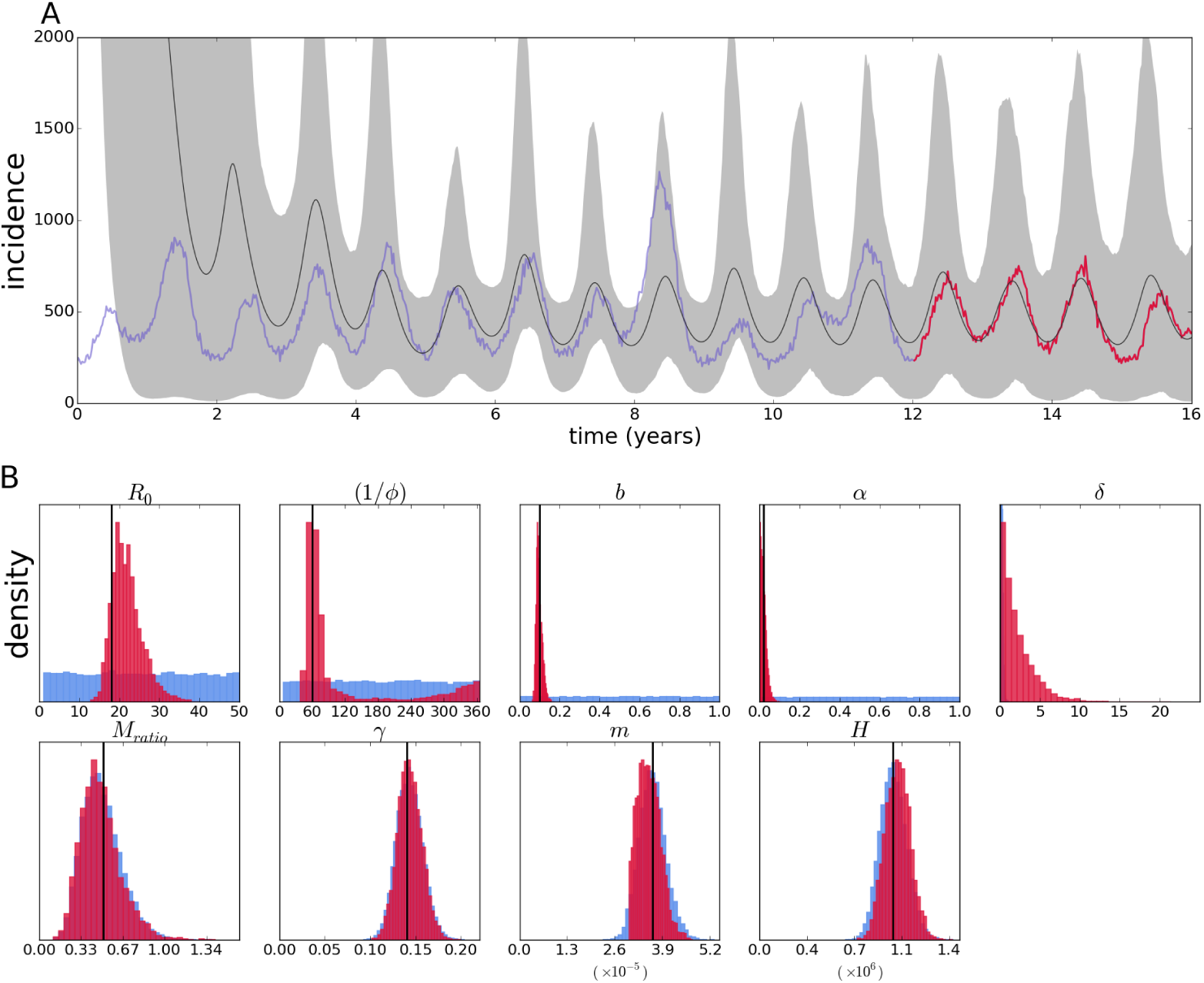
Reconstruction of the epidemic with a two-serotype model from sequences simulated by its corresponding individual-based model (A). Posterior distributions (crimson color) and priors (light blue) of epidemiological parameters (B).

The general *MCMC* settings are the same as with the time series wherever applicable, despite them being different implementations as explained in the methods section. Also, there is no way around the fact that some parameters in this estimation are absent and do not apply to the previous case – for instance the origin time of the epidemic (which for a time series is trivially defined as the first time point), the mutation rate and the tree itself (both of which have no impact on the incidence series). Otherwise, the estimates are generally comparable.

Biases are not nearly as pronounced as in the inference with the time series; Besides, the variables with gamma-distributed priors seem not to get much information from the likelihood and stay almost unaltered compared to their priors. The origin parameter, the time before the present when the epidemics starts, is fixed at 5852 days; this choice is discussed in the following sections. On the other hand, parameters with uninformative priors such as the mosquito mortality rate are quite well estimated; moreover, the basic reproductive number is quite accurately estimated.

#### 3.2.3 Inference for epidemiological data from the city of Rio de Janeiro, Brazil

We apply the methods tested above to epidemiological data from the city of Rio de Janeiro, Brazil. Weekly incidence data for a period of approximately four years between the end of 2009 and that of 2013 was used to fit both the vector SIRx2 and the two-serotype models. Given the caveats observed for the simulated data, preliminary estimation was performed using the *MCMC* method for time series described above. In addition to the continuous model described above, a fixed scale (or reporting rate) parameter multiplies the incidence output to account for underreporting of cases in the recorded series – i.e. a parameter value lower than one means the actual epidemic is larger than the observed time series record.

The parameter estimates for the vector SIRx2 model are shown as posteriors (Figure 14B); the parameters with independent estimates (included in the inference method as prior probabilities) deviate from that expectation, notably the population size *H*, and human mortality rate *m*, both of which are grossly overestimated. It could be that there is a biological explanation for the departures from the independent estimates, but as seen in the previous subsection, bias and correlation between parameters is likely to affect the estimates. Another well accepted value is recovery happening in around a week (1*/γ*); the posterior confidently places that at around twice that time. The biological parameters of interest without accepted values place the mosquito lifespan (1*/b*) at around 22 days, and temporary immunity period (1*/φ*) at 1 day, while *R*_0_ is estimated to be around 8. Parameters of more epidemiological interest are the reporting rate (“scale” parameter) close to only 4% and the vector to human ratio of close to 2 mosquitoes for every human host. The general fit (Figure 14A) is good, although the model predicts a lower incidence at the first and third peaks, as well as an early outbreak around the 20th epidemiological week, which is not present in the epidemiological data.

**Figure 14.**
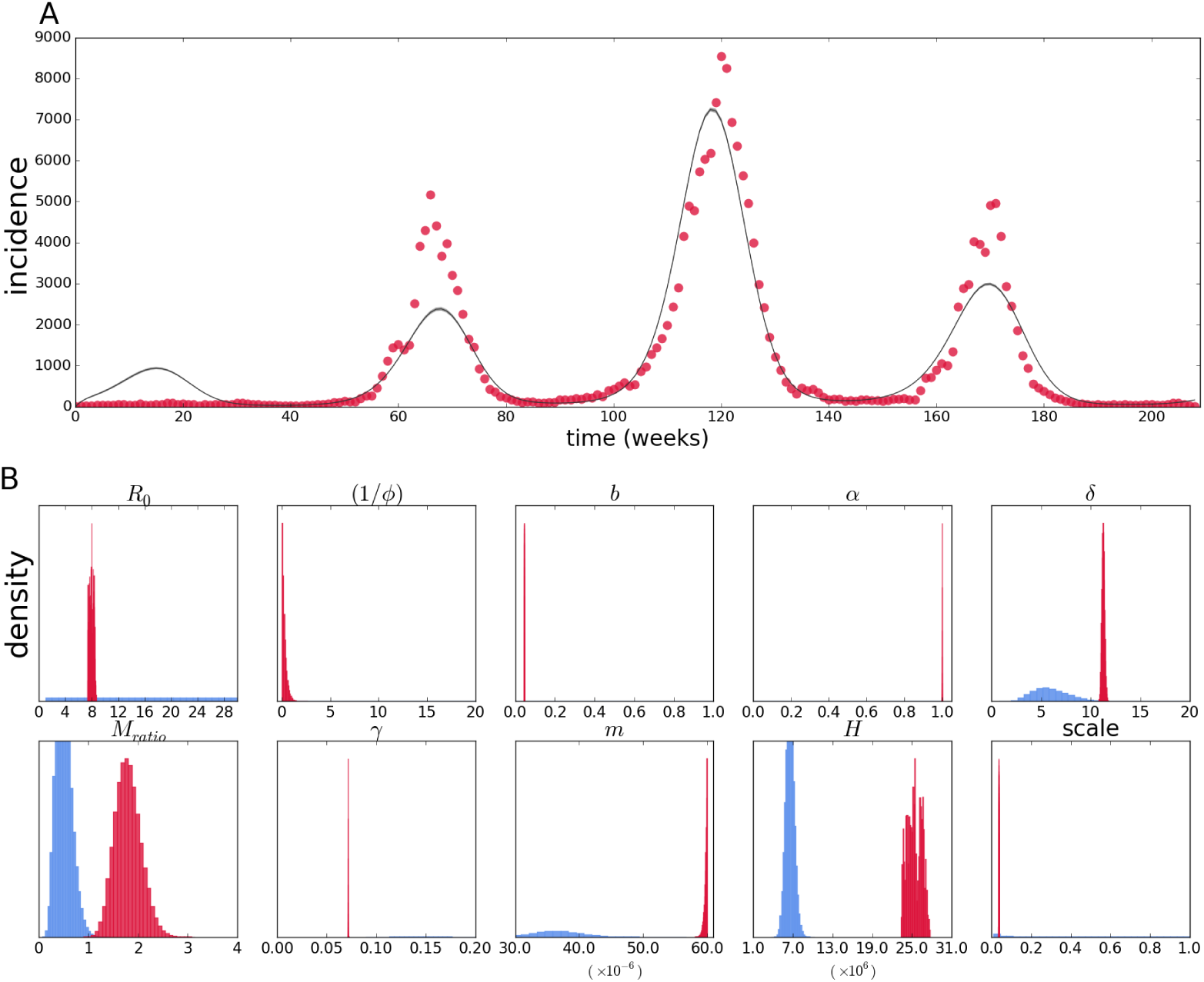
Fit of the vector SIRx2 continuous model with poisson likelihood to epidemiological data from Rio de Janeiro (A). Posterior distributions (crimson color) and priors (light blue) of epidemiological parameters and initial conditions (B).

Next, the two-serotype model is fitted to the same data set; the results are shown in figure 15. The general fit is visibly improved. Interestingly, again some of the better known, or accepted, parameters deviate from expectation in the same direction as with the vector SIRx2 model; human lifespan (1*/m*) is again underestimated, but population size *H* is inferred to be close to the census-recorded number. Recovery *γ* is again estimated to be twice as slow as the commonly accepted rate of a seven day long infection; mosquito mortality rate *b* also has a similar value to that in the one-serotype model.

**Figure 15.**
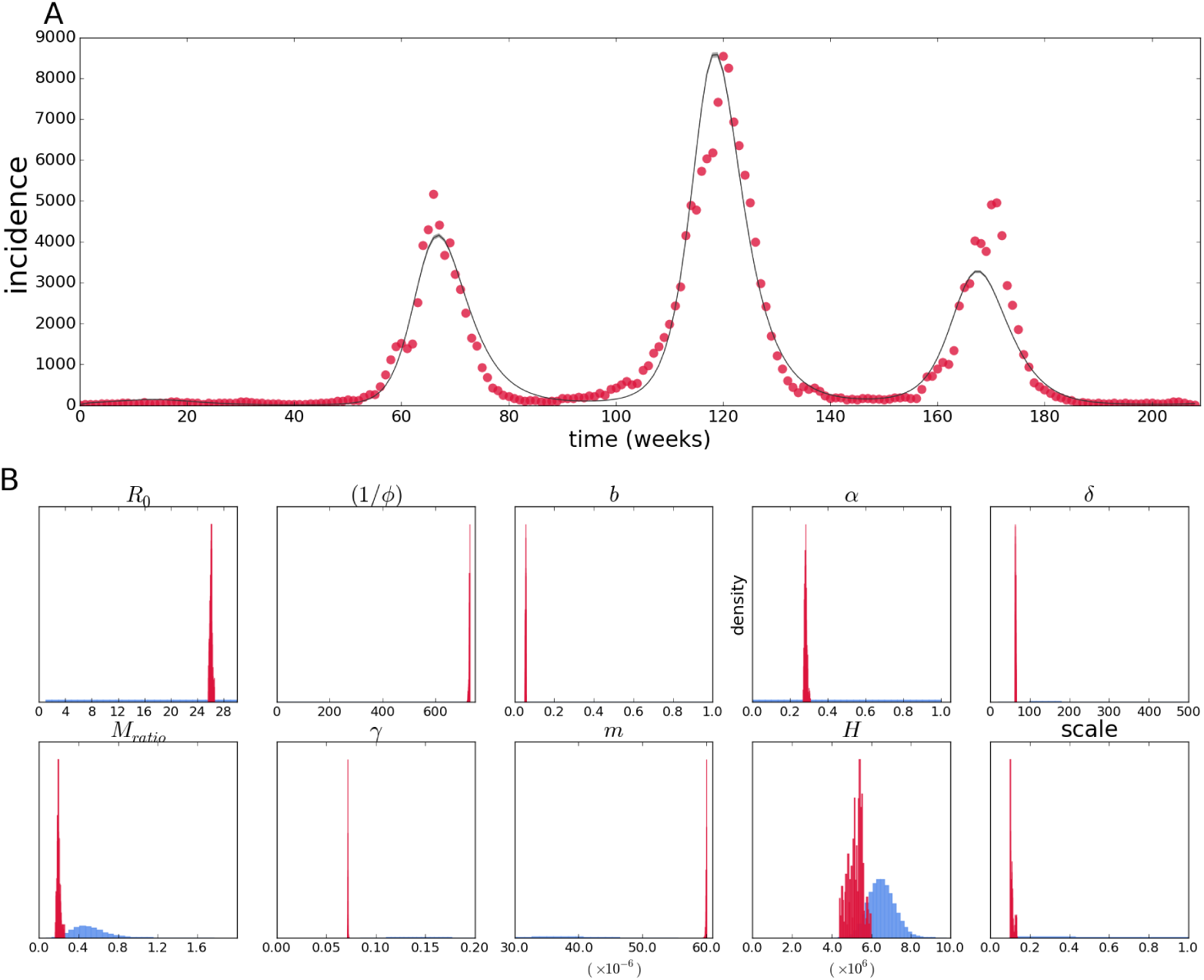
Fit of the two-serotype continuous model with poisson likelihood to epidemiological data from Rio de Janeiro (A). Posterior distributions (crimson color) and priors (light blue) of epidemiological parameters and initial conditions (B).

The robustness of the estimates can be interpreted as a strong signal in the data for these parameters, although it is difficult to know how much they are affected by the deviation in the independently-estimated values, as well as the others with no accepted values such as the temporary immunity period 1*/φ*. The robustness of some of the estimates can be further tested by fixing the parameters with well accepted values (*γ, m, H*). These results are in appendix A. Figure A.6 shows that while preventing parameter values very inconsistent with prior beliefs or independent estimates, most estimates are affected by the choice of free parameters estimated.

Contrary to the vector SIRx2 model, the estimate for *R*_0_ is on the high end of the uniform prior probability assigned to the variable, around 26; the mosquito to human ratio is also opposite to the estimate of the previous model, being about twice as low as the mean of the prior distribution at 0.2, and the cross protection time is in the complete opposite side of the range at 729 days. Mosquito lifespan is close to the previous estimate, 18 days, and the reporting rate is on the order of 10%

The values inferred, however, must not be taken at face value, considering the limitations observed in the inference using the simulated data. We discuss that further in the next section.

## 4 Discussion

The issue of modeling dengue virus transmission is far from trivial, as is probably modeling most infectious diseases. Writing on a paper or programming a SIR-like model into a computer is indeed straightforward; that in its simplicity that alone can capture unintuitive features from disease transmission is quite surprising, but going beyond that is a long and winding road, and extending the basic models in the right direction even more so. It may also be reasonable to ignore variation and selection in some cases (even out of necessity), which greatly simplifies model formulation; still under these simplified models there is plenty to criticize in the existent body of work in modeling dengue transmission (Johansson et al. 2011). Basic simplifications whose consequences are taken for granted may also rear up their ugly heads when the model needs to be confronted to real data.

Inference is a logical next step, although it is an arduous task that may not be as rewarding as straightforward simulation. Despite perfection being far out of sight, the critical exercise can be a constructive one. The difficulties in the case of dengue are illustrated by the issue of multiple serotype interactions, and more subtle effects that are expected – like asymmetry between serotypes or antibody dependent enhancement.

From the forward simulation point of view, it is clear that the structure of the model radically affects the observed incidence – in what can be understood as the crucial factor of the availability of susceptibles – therefore interpreting epidemic series in terms of a single preferred model and its associated parameters is guaranteed to be problematic. The impact of structural differences for a given parameter set – as shown by the forward simulations – is evident, others may not be so clear beforehand, but are indisputable once we become aware of it; for instance, heterogeneity in any rates is more than expected, it must be present, and can have dramatic consequences (Gomes et al. 2012, 2016). These effects are likely to be larger than those of small asymmetries.

A baseline problem of this order is likely not to be solved by adding parameters and processes to the same model structure (Johansson et al. 2011), but instead the basic structures and multiple extensions have to be systematically compared. In theory this problem could be solved by formally comparing the performance of all available models against real data; in practice, the task is a harder one: a record of a time series that aggregates all kinds of infections may lack the information necessary to distinguish between alternative models.

Identifiability analyses may uncover structural features of the model that may prevent inference of particular parameters, and may point to reparametrization of combinations that prevent structural identifiability issues (Bellu et al. 2007); however, it can be cumbersome to implement for larger models (Eisenberg et al. 2013), and it is uncommon (and possibly not feasible) that researchers in the different communities go about all the methods that could improve their results. The analysis of the data simulated by a continuous model strongly suggests that there are no severe structural issues with the method, although I did not perform any of the above-mentioned analyses. Alternative to structural identifiability analyses are more empirical assessments of parameter inferability (Toni et al. 2009), and model comparison based on information criteria (Gelman et al. 2013, chap. 7).

Formal model selection criteria rely on a reasonable fit by the different models; conversely, there should be enough information in the data to grant support to the most appropriate model – for instance, a more or less regular oscillating pattern produced by a two-serotype model may be easily reproduced by a one-serotype model, and the latter may be favored for having fewer parameters; what is more, a single cosine function (or a couple of them) could fit the pattern with fewer of parameters, but that does not change the fact that the data was produced by, and the correct model is still, a two-serotype model.

An example of the difficulties mentioned above are the estimates obtained here from incidence data of the city of Rio de Janeiro. Similar parameter values for different models suggest robustness in the estimates; nevertheless, large deviations from independent estimates may call into question these supposedly robust estimates. Conversely, disparate values for different models may point to the inadequacy of one of the models, and lack of robustness of estimates under model misspecification, but it does not guarantee that either estimate is correct – it is not possible to check deviation from the correct value if no independent estimate is available. Fixing the known parameters can be an additional constraint to the parameter space; however, it may force the remaining parameters into erroneous values due to the decreased flexibility in the model.

The more fine-grained the data is, the more precise can the comparison between models be, but that assumes that some of the models can explain the data well enough. It may, however, be the case that more detailed data causes the models only to fail more miserably than before. In the end, all models are approximations (at best, reasonable ones, but most likely rather crude ones), so that the multiple aspects of model and inference framework must also improve as data becomes better and more plentiful.

The use of sequence data for model-based inference presents itself as both an exciting perspective as well as a challenge: on the one hand it can make important distinctions such as genotypes and serotypes of pathogens, and by design allow inference about the entire population to be derived from a sample. It also carries, by default, information about more than one time series at the same time – i.e. incidence, prevalence and, if applicable, migration (Volz 2012). On the other hand fitting a model to sequences relies on elaborate constructs that are difficult to visualize and evaluate – some of the improvements that come with new kinds of data are therefore not without new issues.

One thing that seems to be unique to epidemiological models is that the data associated to the pathogens can be acquired simultaneously in different formats, so inference with one kind of data may be independently validated by another kind (e.g. sequence-based reconstruction of the epidemic can be compared to incidence data, as shown for pseudo data). Alternatively, these and other kinds of data (serological, vector population data, etc) can be used in combination to improve estimation, if the problem is scarcity of data.

This aspect is probably an important contrast to coalescent-based methods in fields like conservation genetics, where great strides have been made to incorporate processes like recombination (McVean & Cardin 2005), structured environments (as opposed to change in effective population size) (Mazet et al. 2016) and even allow inference from a single recombining genome (Li & Durbin 2011), but where very little validation of alternative models exists, especially with alternative types of data, which are not available for hundreds or thousands of years ago. Coalescent methods in epidemiology offer the opportunity of trying the methods, and validating them with more stringent criteria.

Increasingly it seems that scarcity is not the main problem (Pybus et al. 2013), but rather the difficulty of inferring multi-dimensional parameter sets (or constraining their space enough via independent estimation of individual parameters), and formally comparing full models in a way that makes the estimates actually useful, in addition to more subtle aspects of data collection (Frost et al. 2015).

Although its effects are clear in the individual-based simulations, the issue of stochasticity in the system state was not directly tackled here; therefore, a deterministic model can be forced to fit a different trajectory only by changing its parameters, even if the deviation is caused by chance. Methods such as particle filtering (or Sequential Monte Carlo) allow tracking of the stochastic system state along time (Ionides et al. 2006); these have been incorporated into genealogy-based estimation (Rasmussen et al. 2014a, 2011), potentially solving the issue of the effect of stochasticity in the population immunological states. These methods apply straightforwardly to population immunological states along time, which in the case on inference from time series is directly correspondent to the likelihood; in the coalescent-based estimation that is one of the components of the inference model, but it is not as easy to illustrate genetic drift in a similar way. Implementing methods that account for stochasticity is beyond the scope of this thesis, although the results shown strongly suggest that the simple fact that stochasticity is present can hamper progress in an otherwise simple task.

It can be tempting to focus specifically on the fine-tuning mathematical models, development new inference methods, and on extensive efforts to gather comprehensive data sets, but it is important to take into account how the weaknesses of each of the steps compound into a larger impediment. Concentrating particularly into just one of these (or other even more particular) aspects may prevent a realistic use of data-driven, model-based analysis. I have shown how model structure, assumptions about stochasticity, prior information and data requirements all deserve specific treatments lest the lack thereof introduces or amplifies biases and inaccuracies in the results, and therefore hope to have contributed to the integration of model building, inference frameworks, as well as future efforts to gather epidemiological data of different kinds.

## Acknowledgements

Acknowledgments I would like to thank the supervisor of this work, Gabriela Gomes. I would also like to acknowledge Paulo Campos for sharing code for the individual-based simulation, Flavio Codeço for sharing data and assistance with the time series-based inference, and David Rasmussen for sharing code and further help with the implementation of genealogy-based inference.

The author was a beneficiary of a doctoral grant from the AXA Research Fund.

