## Appendix for "Multiple-serotype models of dengue virus transmission: simulation study and perspectives for the application of inference in epidemiological surveillance"

### Multiple-serotype models of dengue virus transmission: Supplementary text

### A Time series-based inference

#### A.1 One-serotype model inference

The results for the inference with the one-serotype model from simulated data is shown in figure A.1. Biases similar in magnitude are also observed for this simpler model.

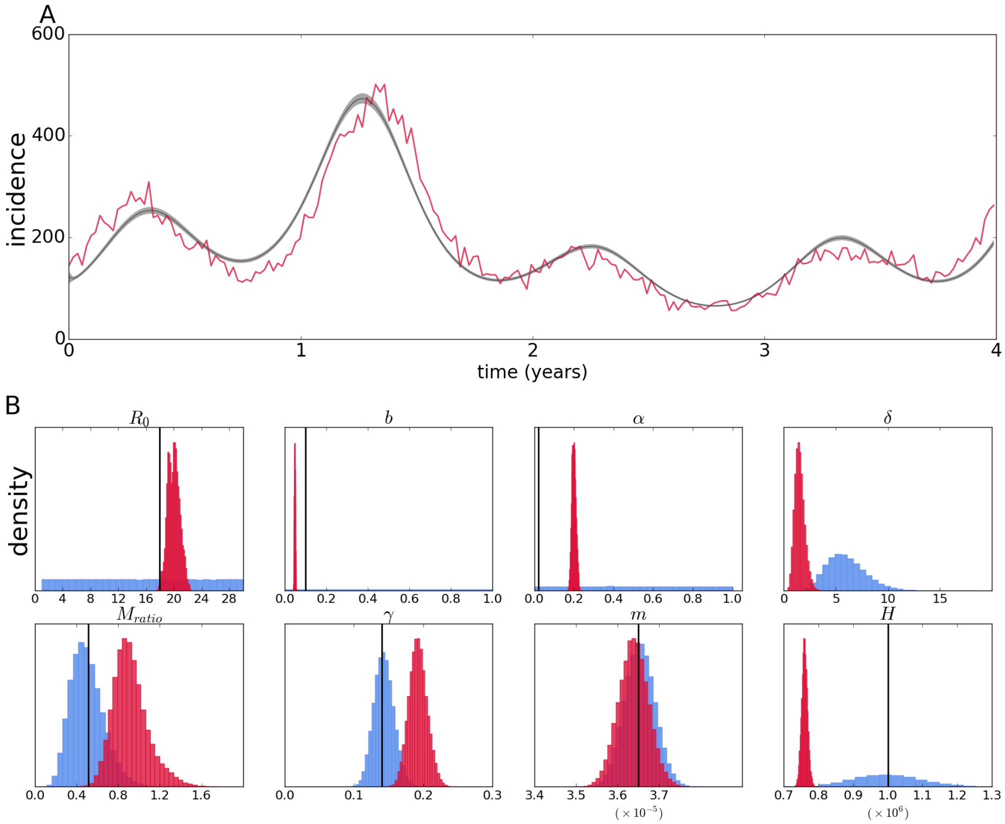


**Figure A.1.** Fit of the one-seroytpe model to stochastically simulated data (A). Posterior distributions (crimson color) and priors (light blue) of epidemiological parameters and initial conditions (B)

#### A.2 Inference of a reduced set of parameters

The inference from time series generated by a continuous model with noise added independently to each time point serves as proof-of-concept for time series as suitable data to infer the parameters in the model; nevertheless, the jump to inference from a fully stochastic model is quite a big one, and testing some intermediate conditions for inference could be warranted. Assuming all epidemiological rate parameters except for $R_0$ were known, the estimates for the parameter are shown in figure A.2, for the case where the initial conditions are inferred and that when these are also assumed to be known.

The bias in the estimate is of the order of around 10% in the first case, and less than 5% when $R_0$ is the only estimated parameter.

Interestingly, when assuming that initial conditions are known, but estimating other parameters for which independent estimates are difficult to obtain, $R_0$ is underestimated by less than 2%.

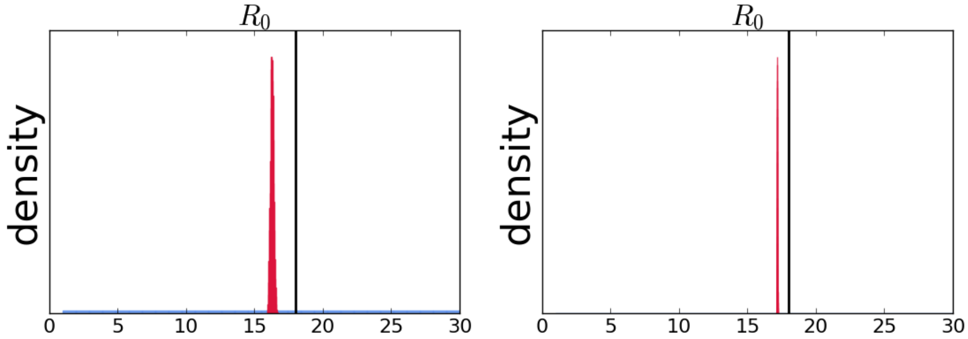


**Figure A.2.** Estimates of $R_0$ as single epidemiological parameter for the case where it is estimated together with the initial conditions of the system (left), or as the only unknown parameter (right)

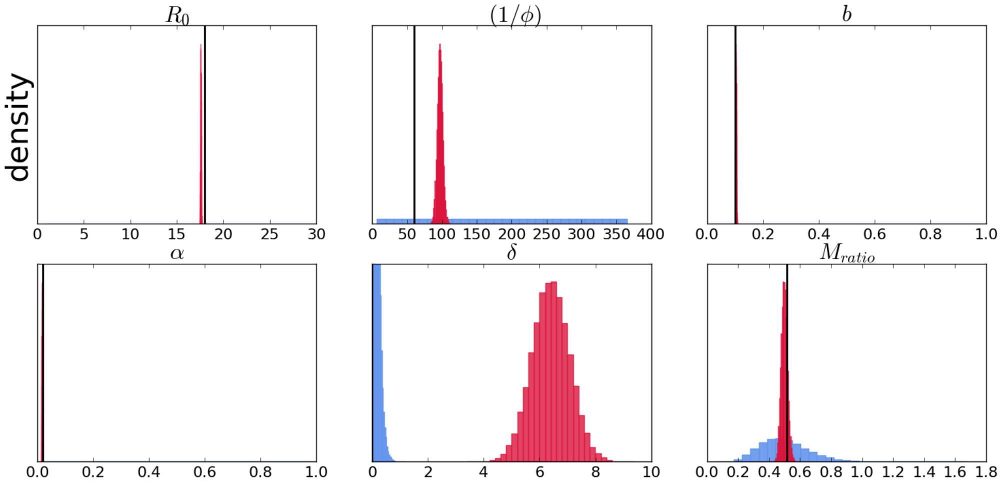


**Figure A.3.** Estimates of a subset of epidemiological parameters assuming initial conditions are known.

These results shown that identifiability and estimation of multiple parameters is an issue, but that the estimation of initial conditions alone can bias the estimation. It is worth noting that even when the known initial values are fixed – not estimated – the time series is still subject to stochasticity, which is not fully accounted by this method or the assumption of known parameters, and can therefore bias the estimates.

#### A.3 Inference under model misspecification

The vector SIR model is a submodel of the vector SIRx2 and the two-serotype model, both of which collapse into it when the immunity waning parameters equals zero (the latter also collapses when only one serotype is circulating). While it is theoretically possible that either model will correctly estimate the appropriate parameters to be zero, in practice the over-specification of the models may affect parameter estimates; conversely, under-specification is guaranteed not to have all parameters, and yet the estimates for the parameters that are still included could be robust to that kind of misspecification.

The result of fitting a one serotype model to a single time series recorded from two-serotypes interacting (but indistinguishable from the data) is shown in figure A.4.

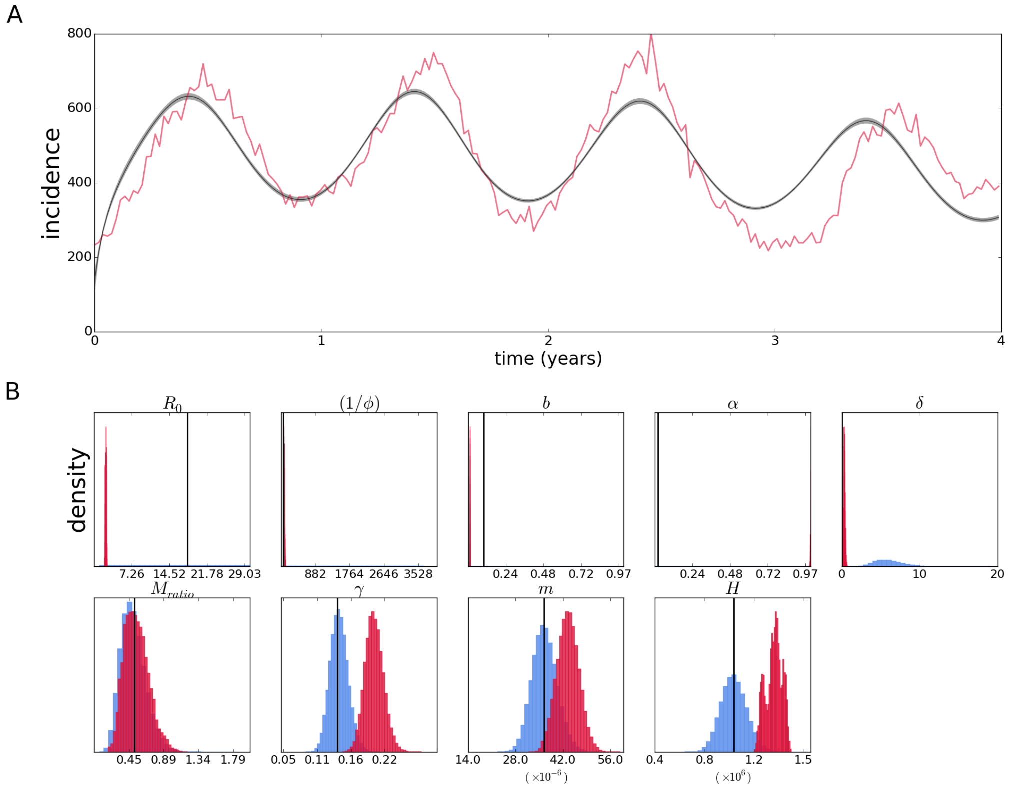


**Figure A.4.** Fit of the vector-SIRx2 continuous model with poisson likelihood to data simulated from a stochastic two-serotype model (A). Posterior distributions (crimson color) and priors (light blue) of epidemiological parameters and initial conditions (B).

Conversely, a model with more than one infected compartment can be fit to data from a single type of infection; that kind over overfitting is shown in figure A.5.

While the fit is visually acceptable, the inference is unable to estimate most parameter values correctly; unexpectedly, convergence seems better against the data simulated by a two-serotype model, although the biases are clear.

This highlights the importance of having enough alternative models to test against the data, and shows that even seemingly convergent estimates may hide significant biases.

#### A.4 Inference of a reduced set of parameters for robustness testing

The results for inference from epidemiological data with the $\gamma$, $m$, and $H$ parameters fixed are shown in figure A.6. The model fit is not visually affected by the fixed parameters. Parameters $\phi$ and $M_{ratio}$ are radically affected; mosquito mortality rate $b$ also changes considerably, while the rest shift more slightly. Particularly, $R_0$ is shifted down to 14.

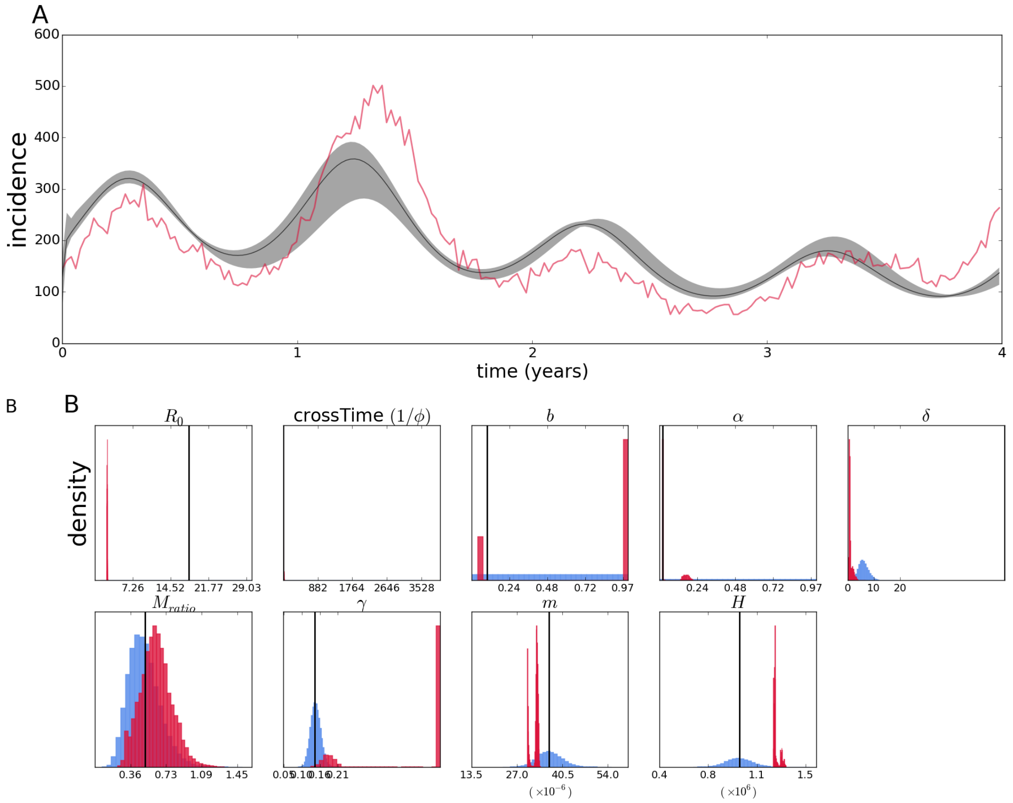


**Figure A.5.** Fit of the vector-SIRx2 continuous model with poisson likelihood to data simulated from a stochastic vector-SIR model (A). Posterior distributions (crimson color) and priors (light blue) of epidemiological parameters and initial conditions (B).

### B Genealogy-based inference

#### B.1 Alternative priors

Considering the effect of stochasticity on the system state, as well as the apparent lack of information about it in the data, it is expected that different priors for these parameters may affect the inference. Arguably the strongest assumption about the initial conditions is that they were completely known, and therefore are kept fixed for estimation of the remaining parameters for the epidemiological rates. If the conditions are observed, or estimated in any manner, they can be treated as less of a certainty and incorporated probabilistically, into the likelihood function in addition to the coalescent process, or time series data.

A weaker imposition is to consider information about the system state as priors in the bayesian method. That too would have different gradations, ranging from strong priors with a mode at a given value, uniform priors over some larger or narrower ranges, or some other sensible choice of priors. In any case, the choice must be justified, but is probably not the strongest assumption embedded into the inference framework. I show the effect of alternative choice of priors for the population initial conditions on the estimates, and detail in which case the priors could be an appropriate choice.

In the case where there is an estimate for the state of every population described by the model, but as mentioned above it is not treated as true, or incorporated into the likelihood, gamma-distributed priors could be used to make the *MCMC* chain tend towards those values unless there is stronger information in the likelihood that brings

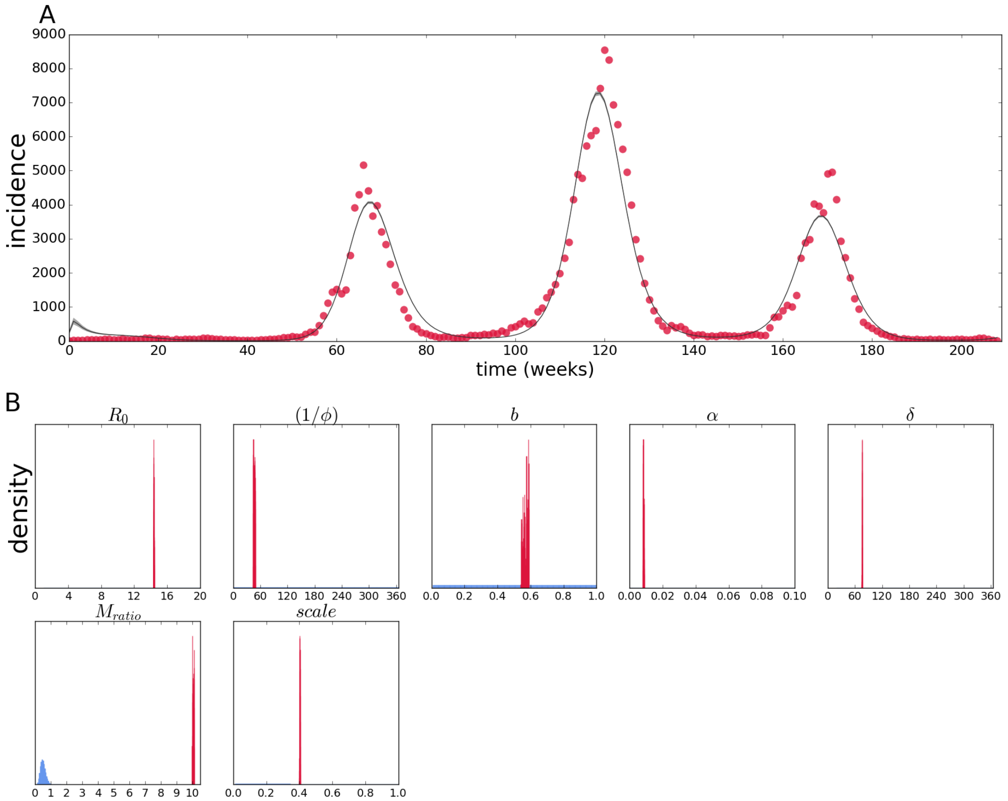


**Figure A.6.** Fit of the two-serotype continuous model with poisson likelihood to epidemiological data from Rio de Janeiro (A), with $\gamma$, $m$, $H$ parameters fixed. Posterior distributions (crimson color) and priors (light blue) of epidemiological parameters and initial conditions (B).

them elsewhere. The results for this case, all else being equal to the main text, are shown in figure B.1. Since there seems not to be a strong signal to estimate the population state parameters when using priors that are uniform or nearly so, using these parameters will essentially recover the prior distribution. The estimate for $R_0$ is more precise, although $1/\phi$ is biased to lower values, and $\alpha$ is less precise. Otherwise, the difference is not striking.

The opposite of the above is using uniform priors over an extremely range of possible values (since here we know that the total population is a million people, and that that is enforced by the gamma-distributed prior on that parameter, this is roughly translated to uniform over the range from zero to a million, which could also be made even larger for the sake of it). For these wide uniform priors, the results are shown in figure B.2.

For the human infected compartments shown in panel C of figure B.2, instead of having the posteriors squeezed close to zero, they are distributed over a wider interval, although the susceptible and recovered parameters do not differ notably from what was observed before. The most striking difference is the vector infected compartments are estimated to have values on the higher extreme of the distribution, which in this case is known to be wrong, as both human and vector infections oscillate in around a few thousand of infections at any point in time. As for the epidemiological parameters, shown in panel B, $R_0$ is overestimated in this case, and $1/\phi$ is overestimated with very little precision, covering most of the uniform prior (with notable exception of the actual value). Nevertheless, reconstruction of the epidemic is not severely affected, but does

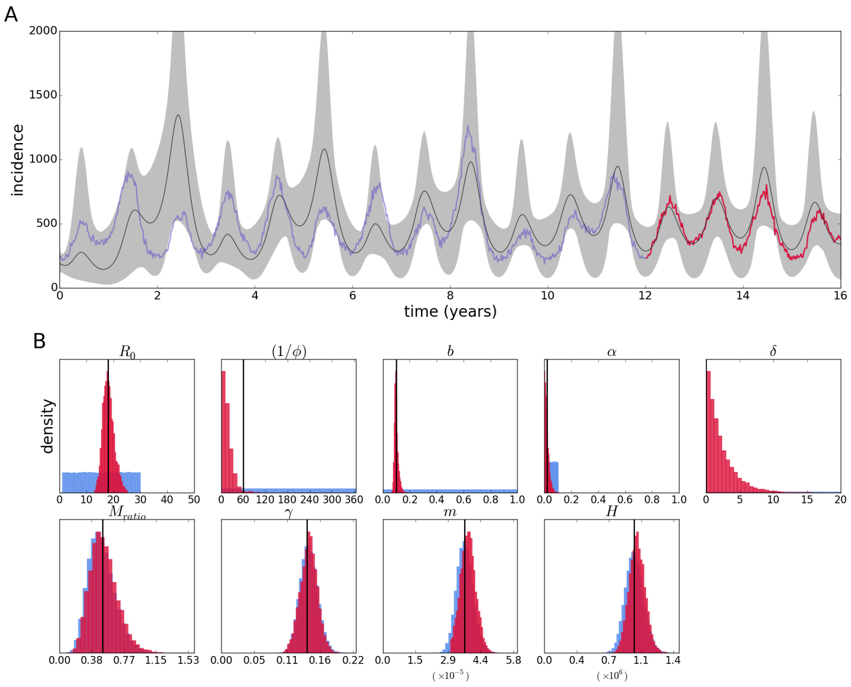


**Figure B.1.** Reconstruction of the epidemic (A), and posterior distributions (crimson color) and priors (light blue) of epidemiological parameters (B) for the two-serotype model and gamma-distributed priors for the initial conditions.

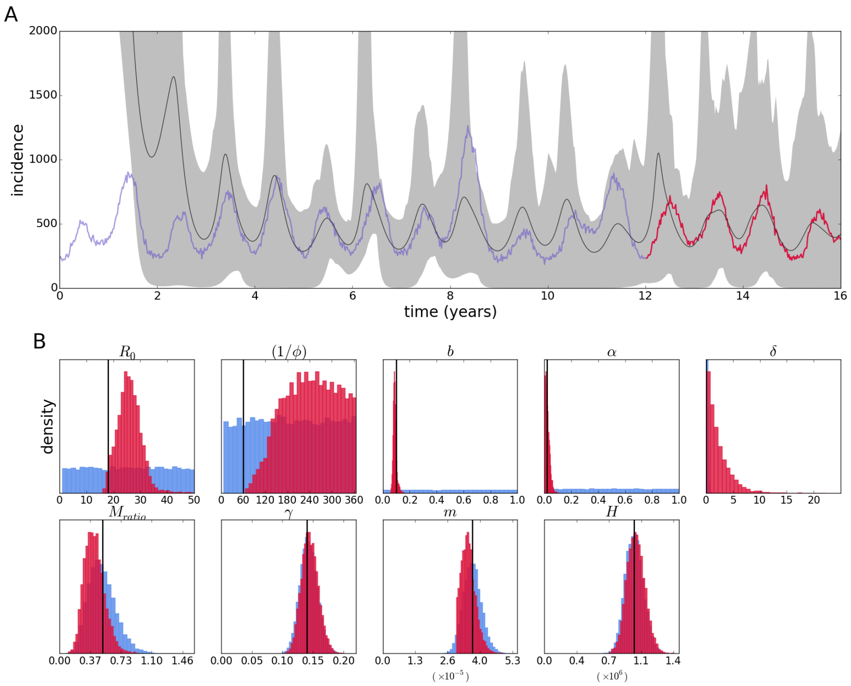


**Figure B.2.** Reconstruction of the epidemic (A), and posterior distributions (crimson color) and priors (light blue) of epidemiological parameters (B) for the two-serotype model and wide uniform priors for the initial conditions.

have wider credibility intervals.

A less conservative use of uniform priors is that with different ranges depending on the populations. Because the human infected compartments are at least partially observed, it may be reasonable to define a narrower uniform distribution for these variables More generally it is easy to do that for the simulated data because it is known what is a reasonable range for each variable; nevertheless, when that is unknown or doubtful there is the risk of limiting the interval too much and excluding the correct range. The results for this specification for the priors is shown in figure B.3. The

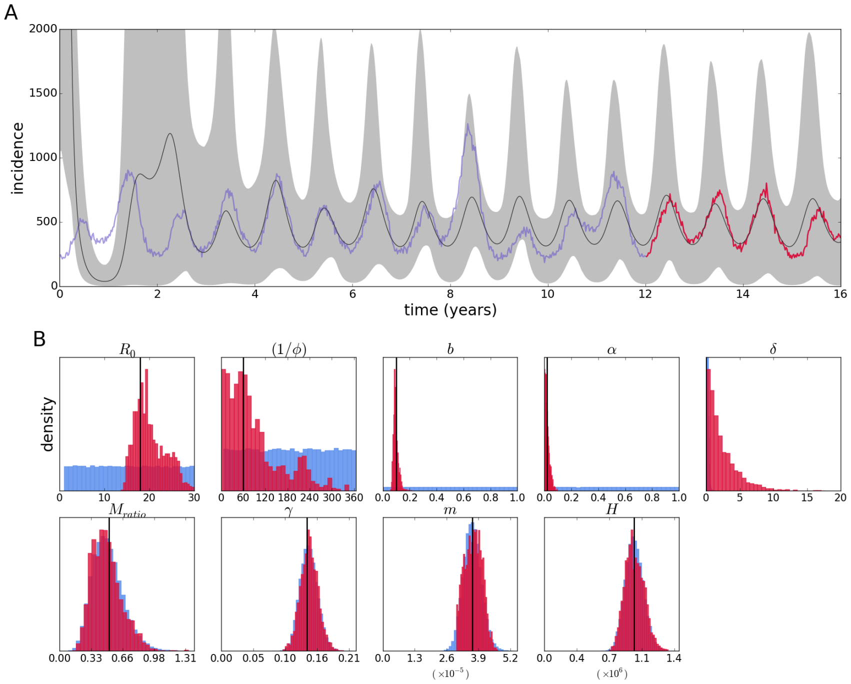


**Figure B.3.** Reconstruction of the epidemic (A), and posterior distributions (crimson color) and priors (light blue) of epidemiological parameters (B) for the two-serotype model and wide uniform priors for the initial conditions.

estimates of the epidemiological parameters (figure B.3, panel B) are similar to those obtained before, with somewhat less precision in the estimate of $R_0$ compared to the results in the main text, or the gamma-distributed priors, but the posterior for $1/\phi$ includes the true value better the the latter - although the distribution is quite wide. Regarding the population states, the posteriors are not very precise compared to the priors, but even that can mean a better precision than the posteriors obtained with wide uniform priors, since the latter can span 3 or 4 orders of magnitude more. The reconstruction of the epidemic has somewhat narrower credibility intervals, and overall inference is improved compared to the wide priors.

#### B.2 One serotype subset for two-serotype inference

Furthermore, inference for the two-serotype model can be made by computing the likelihood of the coalescent process for only one serotype, so results are shown for that case in conditions similar to that of the two serotypes. The results for gamma-distributed priores are shown in figure B.4. For completely flat priors, using the subset of the data corresponding to the second seroytpe, the results are shown in figure B.5. For the counterpart of the data set, that is the first serotype, the estimates are

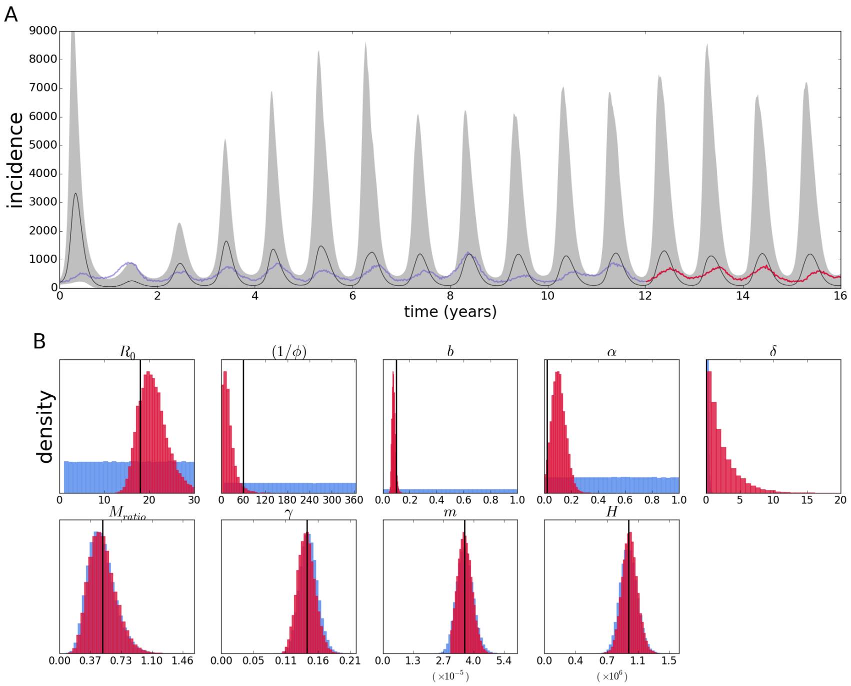


**Figure B.4.** Reconstruction of the epidemic (A), and posterior distributions (crimson color) and priors (light blue) of epidemiological parameters (B) for the two-serotype model and gamma-distributed priors for the initial conditions, using data from only one of the serotypes ("serotype 2").

shown in figure B.6. In this case, the estimates can be seen to be worse than the alternative serotype; because the amount of data is comparable, the difference could in principle be attributed to stochasticity in the mutations that happened to appear and were sampled in each data set.

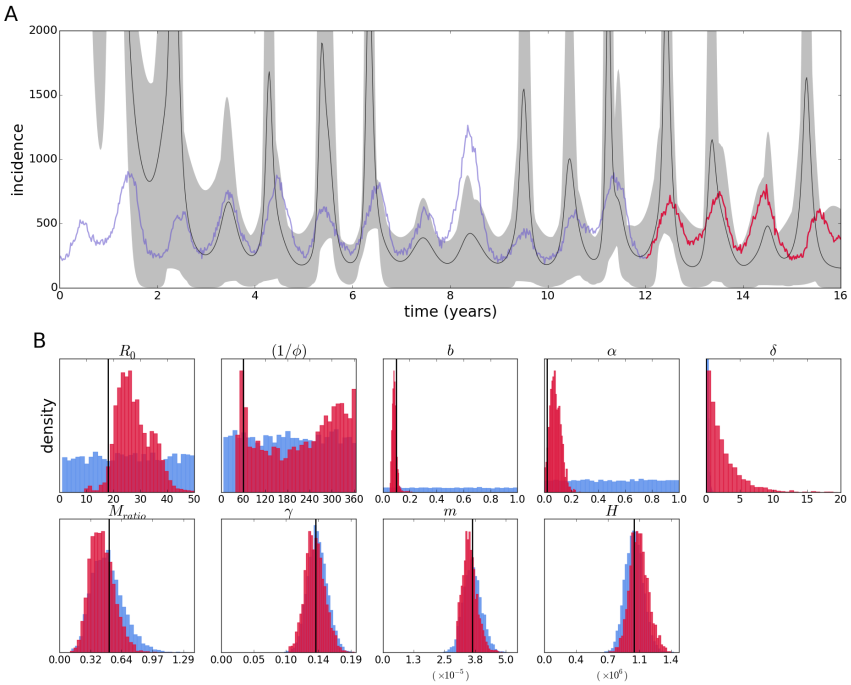


**Figure B.5.** Reconstruction of the epidemic (A), and posterior distributions (crimson color) and priors (light blue) of epidemiological parameters (B) for the two-serotype model and wide uniform priors for the initial conditions, using data from only one of the serotypes ("serotype 2").

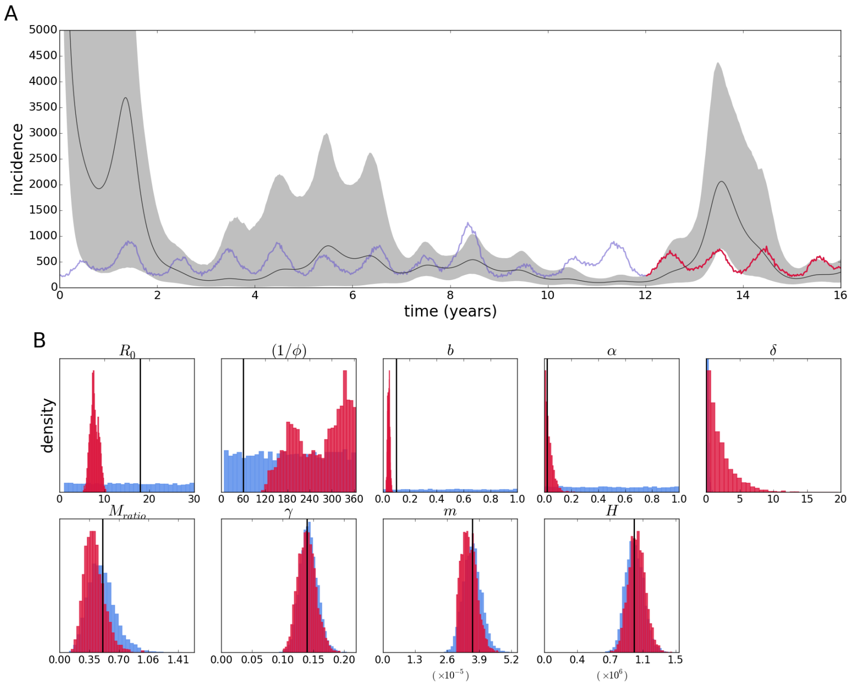


**Figure B.6.** Reconstruction of the epidemic (A), and posterior distributions (crimson color) and priors (light blue) of epidemiological parameters (B) for the two-serotype model and wide uniform priors for the initial conditions, using data from only one of the serotypes ("serotype 1").
